# Enhanced autophagic-lysosomal activity and increased BAG3-mediated selective macroautophagy as adaptive response of neuronal cells to chronic oxidative stress

**DOI:** 10.1101/580977

**Authors:** Debapriya Chakraborty, Vanessa Felzen, Christof Hiebel, Elisabeth Stürner, Natarajan Perumal, Caroline Manicam, Elisabeth Sehn, Franz Grus, Uwe Wolfrum, Christian Behl

## Abstract

Oxidative stress and a disturbed cellular protein homeostasis (proteostasis) belong to the most important hallmarks of aging and of neurodegenerative disorders. The proteasomal and autophagic-lysosomal degradation pathways are key measures to maintain proteostasis. Here, we report that hippocampal cells selected for full adaptation and resistance to oxidative stress induced by hydrogen peroxide (oxidative stress-resistant cells, OxSR cells) showed a massive increase in the expression of components of the cellular autophagic-lysosomal network and a significantly higher overall autophagic activity. A comparative expression analysis revealed that distinct key regulators of autophagy are upregulated in OxSR cells. The observed adaptive autophagic response was found to be independent of the upstream autophagy regulator mTOR but is accompanied by a significant upregulation of further downstream components of the canonical autophagy network such as Beclin1, WIPI1 and the transmembrane ATG9 proteins. Interestingly, the expression of the HSP70 co-chaperone BAG3, mediator of *BAG3-mediated selective macroautophagy* and highly relevant for the clearance of aggregated proteins in cells was found to be increased in OxSR cells that were consequently able to effectively overcome proteotoxic stress. Overexpression of BAG3 in oxidative stress-sensitive HT22 wildtype cells partly established the vesicular phenotype and the enhanced autophagic flux seen in OxSR cells suggesting that BAG3 takes over a key part in the adaptation process. A full proteome analysis demonstrated additional changes in the expression of mitochondrial proteins, metabolic enzymes and different pathway regulators in OxSR cells as consequence of the adaptation to oxidative stress in addition to autophagy-related proteins. Taken together, this analysis revealed a wide variety of pathways and players that act as adaptive response to chronic redox stress in neuronal cells.

## 1. Introduction

Oxidative stress, the accumulation of reactive oxygen species which chemically modify and damage proteins, lipids and DNA disturbs cellular homeostasis and has been linked to aging and the pathogenesis of diseases such as cancer and neurodegeneration [1]. Neuronal disorders like Alzheimer’s Disease (AD), Parkinson’s Disease and Amyotrophic Lateral Sclerosis (ALS) show increased levels of oxidation end products in affected brain regions [2], [3]. Strikingly, in AD nerve cell death is not equally distributed throughout the brain but it is rather occurring in the pyramidal neurons of the entorhinal and the inferior temporal cortex, the hippocampus and the amygdala while other brain regions such as the cerebellum show resistance [2], [4], [5]. Some aspects characterizing the basis for this selective vulnerability of neurons in distinct brain regions were addressed before and pinned down to the ability to actually resist oxidative stress [6]. Also, in other neurodegeneration pathologies selective vulnerability (and resistance) is observed [7]. A highly upregulated antioxidant defense is also linked to the progression of certain tumor types [8], [9]. Though oxidative stress resistance appears to be disadvantageous in the context of tumor therapy, it could be beneficial in the attempt to support long-term survival of cells that are particularly sensitive to oxidative stress, such as post-mitotic neuronal cells. Therefore, it would be a huge step forward to uncover the molecular basis of an adaptation to oxidative stress and redox stress-associated pathological mechanisms.

To overcome challenges of the cellular homeostasis network during elevated cellular stress, autophagy plays a crucial role. In this process, double-membraned vesicles, called autophagosomes, are formed which sequester large protein complexes and organelles and finally deliver them to lysosomes for degradation [10]. Functionally, autophagy is an intracellular recycling process that guarantees energy supply by generating amino acid building blocks during low nutritional condition [11]. Additionally, autophagy works as an essential stress-adaptive response and rescue mechanism by selective degradation of damaged organelles, protein aggregates and intracellular pathogens [12], [13]. Recently, autophagy was found to be a survival response against oxidative stress in bone marrow–derived mesenchymal stromal cells [14] and, moreover, the activation of autophagy can protect against oxidative stress-induced neuronal damage in adult rats [15].

In the context of autophagy and oxidative stress, previously it has been shown that the HSP70 co-chaperone Bcl2-associated athanogene 3 (BAG3) is highly upregulated in many aggressive tumor types and a link between oxidative stress, enhanced autophagy and BAG3 expression is described for several tumor cells [16] [17], [18], [19]. BAG3 was identified as a key player in the *BAG3-mediated selective macroautophagy* [20] and established as an important partner of the cellular proteostasis network under oxidative and proteotoxic stress as well as in aging conditions [21], [22], [23], [24].

The concept of oxidative stress adaptation has been successfully applied by different groups employing clonal neuronal cells lines, such as rat pheochromocytoma PC12 and mouse clonal hippocampal HT22 cells [25], [26], [27], [28], [29]. Previous studies mainly focusing on the redox stress-resistance phenotype and its reversal in PC12 and HT22 cells revealed key roles for the transcription factor NF-κB, sphingolipids and increased levels of antioxidant enzymes to provide the oxidative stress resistance phenotype [26], [27], [28]. In our current study we now systematically analyzed molecular and functional changes in HT22 cells stably adapted to redox stress as induced by hydrogen peroxide (here called OxSR cells) with a particular focus on the autophagy network. We observed an increased autophagic-lysosomal and a decreased proteasomal activity in OxSR cells and analyzed in detail the expression patterns of key autophagy regulators. We identified that the expression of BAG3 and *BAG3-mediated selective macroautophagy* is upregulated suggesting BAG3 thus may play a particular role in oxidative stress adapted-cells. Finally, a whole proteome comparison between wild type and OxSR cells revealed a wide range of alterations of key proteins involved in different cellular pathways in addition to the autophagy regulators demonstrating the massive impact of chronic redox stress on the protein expression pattern during oxidative stress adaptation.

## 2. Material & methods

### 2.1 Cell culture

Wild type HT22 cell line (HT22-WT), a cloned mouse hippocampal neuronal cell line which is very susceptible to oxidative stress [28], [30] was used as control cell line. HT22 cells resistant to hydrogen peroxide induced oxidative stress, here called OxSR cells, were established by clonal selection. The details of the selection procedure have been described elsewhere [31]. Both cell lines were cultured in Dulbecco’s modified Eagle’s medium containing 10 % fetal calf serum (FCS), 1 mM sodium pyruvate and 1x penicillin/streptomycin (Invitrogen, Karlsruhe, Germany). To maintain the resistant phenotype, 450 µM of H2O2 f.c. (Sigma, Deisenhofen, Germany) was added twice a week to the OxSR cells. Prior to performing experiments, OxSR cells were cultured for three days without H2O2 and medium was exchanged daily to remove residual toxins. Although oxidative stress-resistant mouse hippocampal HT22 cells have been employed before, for the present study we initially reconfirmed the previously observed characteristics of the cell clones used here. So, the cell proliferation rates of the different cell clones were estimated by MTT assay. Consistent with previous findings [31] the growth rate of the OxSR cells was found to be lower than that of the HT22-WT cells (supplementary figure S1A) confirming that increased vitality and oxidative stress resistance of the selected clones was not simply based on a higher proliferation rate.

### 2.2 Pharmacological agents and antibodies

Stock solutions of Bafilomycin A1 (LC Laboratories, B-1080), MG132 (Calbiochem, 474790) and Rapamycin (Enzo, BML-A275-0025) were prepared in DMSO (Roth, A994.2). Stock solution of Canavanine (Santa Cruz Biotech, sc-202983A) was prepared in distilled H2O.

Antibody sources were as follows: for BAG1 (Abcam, ab7976), BAG3 (Proteintech Group, 10599-1-AP), BECN1 (Cellsignaling, 3495), CTSD (Abcam, ab75852), DLP1 (BD Transduction Laboratories,611113), LAMP2 (DSHB Biology, ABL-93), LC3B (Nanotools, 0260-100), LC3B (Sigma, L7543), Phospho mTOR (Abcam, ab109268), mTOR (Calbiochem, OP97), p62 (Progen, GP62-C), PIK3C3 (Cellsignaling, 4263), Poly-Ubiquitin (Dako, Z0458), RAB18 (Sigma, SAB4200173), Tubulin (Millipore, MAB1637), Tubulin (Sigma, T9026), TFEB (Proteintech Group, 13372-1-AP), Vimentin (SCBT, sc-373717), WIPI1 (Sigma, HPA007493).

### 2.3 Plasmids, siRNAs and transfection method

Expression plasmid for mouse FLAG tagged BAG3 (pFLAG-BAG3) was constructed by cloning partial mouse BAG3 cDNA containing the whole CDS into pEGFP-N1 (Clontech). Forward primer sequence used to clone BAG3 plasmid is 5′-AAAGGATCCAGCGCCGCCACCC-3′ and the reverse primer sequence is 5′-GACTCTAGATCACTAGGGAGCCACCAGGTTGC-3′. After vector linearization with BamHI and XbaI, PCR products were inserted using the In-Fusion reaction according to manufacturer’s protocol (Clontech).

Two independent sets of siRNA against BAG3 were purchased from Eurofins MWG Operon with 3′-dTdT overhangs. Sequence of used siRNA1 is 5′-AAUACCUGAUGAUCG AAGA-3′ and siRNA2 is 5′-AUACCUGAUGAUCGAAGAG-3′. Generally, cells were transfected by mixing 20 µg of each of the siRNA as duplex. In all knockdown experiments, same amounts of siRNA targeting a nonsense sequence (5′-AUUCUCCGAACGUGUCACG-3′) were transfected as control.

Cells were transfected by electroporation using the Amaxa Nucleofector I (program U-24) and standard electroporation cuvettes (Sigma). Electroporation buffer details are mentioned elsewhere [21].

### 2.4 Immunoblotting and immunocytochemistry

Western blot analysis and inmmunocytochemistry were carried out as previously described [21]. Blots were developed with the Fusion-SL 3500 WL system (Peqlab, Erlangen, Germany) and densitometry analysis was performed using Aida Image Analyzer v.4.26 software (Raytest, Straubenhardt, Germany). For the labelling of the lysosomes and acidic vacuoles, the fluorescent dye LysoTracker was used. Cells (cultured on 24-well plates with coverslips) were incubated with 100 nM LysoTracker Red DND-99 (Thermo scientific, L7528) in culture medium for 1 h under incubator conditions and then fixed with 4 % PFA. Cells for immunocytochemistry were analyzed using a confocal laser-scanning microscope (LSM710, Zeiss) and images were processed with Adobe Photoshop CS5 (San Jose, CA, USA).

### 2.5 Transmission electron microscopy

Sample preparation and transmission electron microscopy (TEM) analysis were carried out as previously described [21] and images were processed with Adobe Photoshop CS5.

### 2.6 Proteomic analysis

Proteomic analysis was executed in different distinct steps such as protein extraction, sample preparation, 1-dimensional gel electrophoresis (1DE), mass spectrometry (MS) based discovery proteomics analysis, bioinformatics and functional annotation and pathways analyses. General approaches used for the analysis were adapted from previously published papers [32], [33], [34], [35], [36], [37], [38], [39]. Details of the individual techniques employed for this paper are described in supplementary file S3 “proteomics methods”.

### 2.7 Quantitative real time PCR array

RNA extraction, cDNA synthesis and Quantitative real-time PCR (qPCR) for HT22-WT and OxSR cells were performed by the protocol previously described [40]. A commercial ready to use primer mix in 96 well format (Biomol, MATPL-1) containing total 88 different genes involved in autophagy were used for qPCR array. PCR cycles were set according to the manufacturer’s protocol. CT values obtained from Quantitative Real-Time PCR data were applied to REST software for calculating the relative fold change in expression of each gene [41]. Actin was used as a reference gene. A fold change of +1.5 or −0.6 is considered as regulated gene.

### 2.8 Measurement of proteasome activity

This procedure is adapted from previous publication [42]. Briefly, cells were washed with ice cold PBS and lysed in lysis buffer (50 mM HEPES, pH 7.8, 10 mM NaCl, 1.5 mM MgCl2, 1 mM EDTA, 1 mM EGTA, 250 mM sucrose, and 5 mM DTT) at 4°C. The homogenates were centrifuged at 10 000 *g* for 15 min at 4°C. Supernatants were collected and normalized to protein content. Enzymatic reaction was started by mixing active cell extracts (containing 15 μg protein determined by BCA) with 100 μl of assay buffer. Assay buffer composition is same as lysis buffer only supplemented with ATP (2 mM) and proteasome substrate (100 μM, Suc-LLVY-AMC, Sigma). AMC fluorescence released by cleavage of substrate by proteolytic activity was recorded in a black 96-well plate after 60 min incubation at 37°C using the Victor 3V Multilabel counter (Perkin Elmer). Specific proteasomal activity was determined by subtracting unspecific AMC fluorescence obtained in the presence of proteasome inhibitor MG132 (20 μM). Results are expressed as % AMC release.

### 2.9 Cell proliferation assay

The growth rate of HT22-WT and OxSR cells were determined by the analysis of cellular metabolic activity. The ability of cells to reduce yellow coloured water soluble MTT to a blue violet coloured water insoluble formazan crystals were measured colorimetrically over nine alternate days. The detailed procedure is described elsewhere [31].

### 2.10 Statistical analysis

Statistical significance of all quantified Western blots, lysotracker staining and qPCR data was determined by unpaired t-test using GraphPad Prism 7 (GraphPad Inc., San Diego, USA). Growth curve for cell proliferation assay was analyzed by two-way ANOVA with the same software. Statistical significance was accepted at a level of p<0.05. The results are expressed as mean ± S.E.M. The proteomics data were statistically analyzed with Perseus software (version 1.6.1.0). The degree of variances between the biological replicates and groups were assessed by Pearson’s correlation analysis and Student’s two-sided t-test was used for the groups’ comparison to identify the significantly differentially abundant proteins. Details are maintained in supplementary file S3 “proteomics methods”.

## 3. Results

### 3.1 Autophagosomes, autophagic flux and lysosomal activity are increased in OxSR cells

Based on previous findings that HT22 cells resistant to H2O2 (OxSR) show an upregulation of the lysosomal marker LAMP1 compared to oxidative stress-sensitive HT22 wildtype cells (HT22-WT) [28], we analyzed the cellular vesicular ultrastructure in detail. TEM revealed the presence of an intensive vesicular network in OxSR cells and a higher number and size of double-membrane autophagosomes as well as lysosomes in OxSR cells, suggesting an enhanced autophagic-lysosomal activity (Figure 1A). Therefore, we next investigated the baseline autophagic flux in OxSR cells in presence of BafA1 which inhibits fusion of autophagosomes and lysosomes. We observed a significantly elevated autophagic flux, the enhanced accumulation of LC3B-II as well as p62 upon BafA1 treatment in OxSR cells compared to HT22-WT cells (Figure 1B-D). Consistently, immunostainings showed a generally upregulated lysosomal activity in OxSR cells indicated by the lysosomal enzyme marker cathepsin D (Figure 1E). Nuclear translocation of TFEB is known to have a pivotal role in upregulation of autophagy and lysosomal biogenesis [43]. TFEB was found to be localized mainly in the cytoplasm of HT22-WT cells and almost exclusively translocated to the nucleus in OxSR cells as shown by immunocytochemistry (Supplementary Figure S1B) pointing towards an ongoing transcription of TFEB target genes in OxSR cells supporting the continuous autophagic activity.

**Figure 1:**
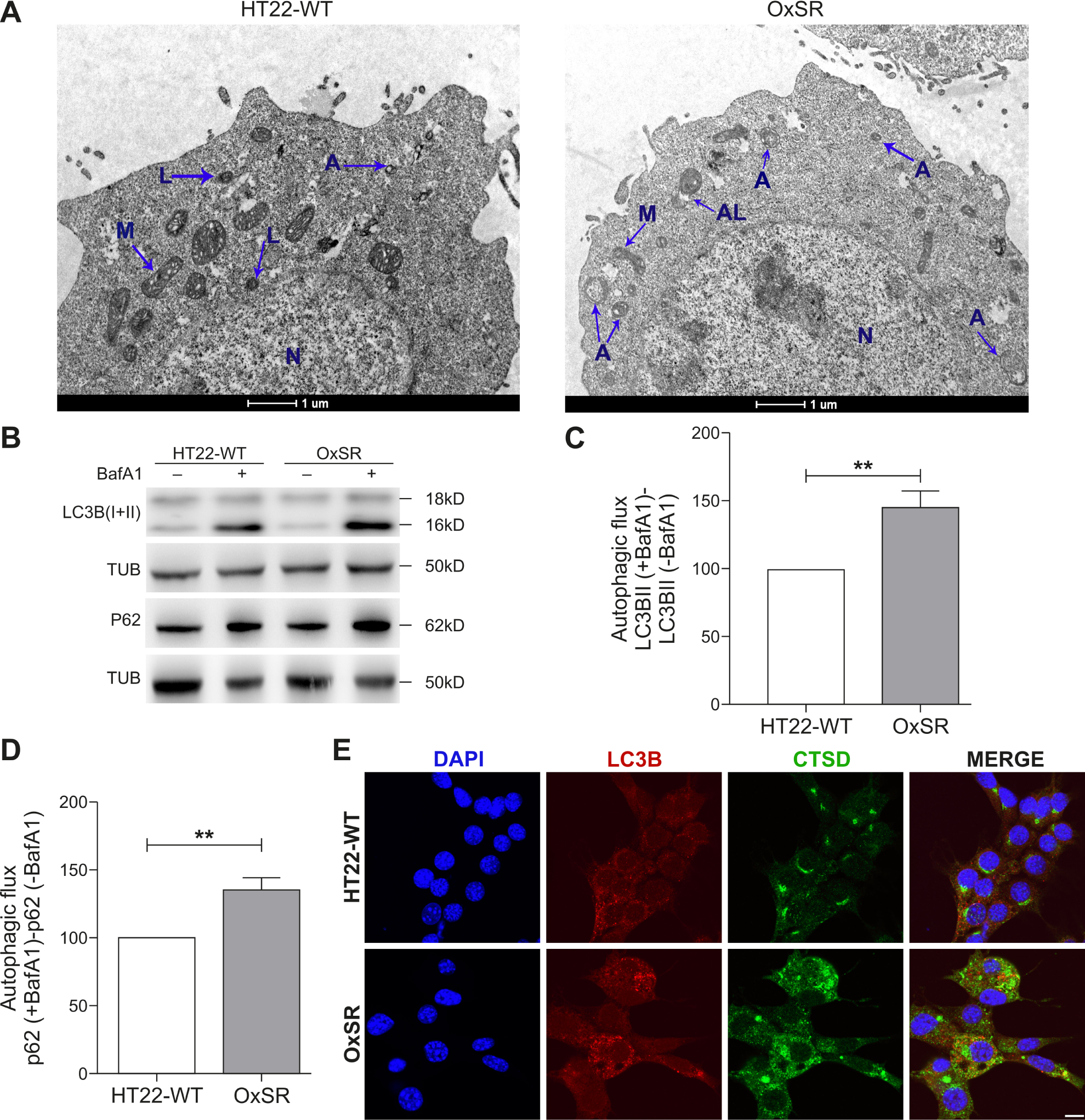
OxSR cells show higher autophagic activity. (A) Ultrastructure analysis of HT22-WT and OxSR cells by TEM. OxSR cells clearly showed increased number and size of highly electron dense vacuolar autophagic vesicles at different stages of maturity (representative images are shown). Autophagosomes are indicated by A, autolysosomes by AL, nucleus by N and mitochondria by M. (B) Protein extracts from HT22-WT and OxSR cells have been taken after 6 h of BafA1 (8 µM) treatment and expression of the indicated proteins was detected by Western blotting. (C, D) The autophagic flux was measured after densitometric analysis. Therefore, tubulin normalized LC3B-II and P62 levels in the absence of the lysosomal inhibitor were subtracted from corresponding levels obtained in the presence of BafA1. Tubulin (TUB) is used as loading control. Values represent mean ± S.E.M., n=3, **p<0.01 and control HT22-WT cells were set to 100%. (E) HT22-WT and OxSR cells were immunohistochemically stained against the LC3B and Cathepsin D (CTSD). Scale bar: 10 µm. Pictures were taken by confocal microscopy.

### 3.2 OxSR cells show enhanced expression of several key autophagy regulators but mTOR expression and phosphorylation status is unchanged

Next, we focused on the expression of certain key regulators of canonical autophagy in OxSR cells (as indicated in the schematic diagram in Figure 2A). Employing Western blotting no changes in the phosphorylation status of the main regulator of canonical autophagy mTOR (ratio of p-mTOR to mTOR as measure of mTOR-dependent autophagy induction) was detectable in the OxSR cells (Figure 2B and C). To check the responsiveness of the mTOR-regulated autophagy pathway in OxSR cells, we treated the cells with rapamycin and monitored autophagic flux after co-treatment with BafA1. We found that rapamycin significantly increases autophagic flux in OxSR cells indicating that the mTOR-mediated autophagic pathway is still principally inducible in this cell line (Figure 2H and I) but obviously is not activated under basal conditions in OxSR cells adapted to redox stress.

**Figure 2:**
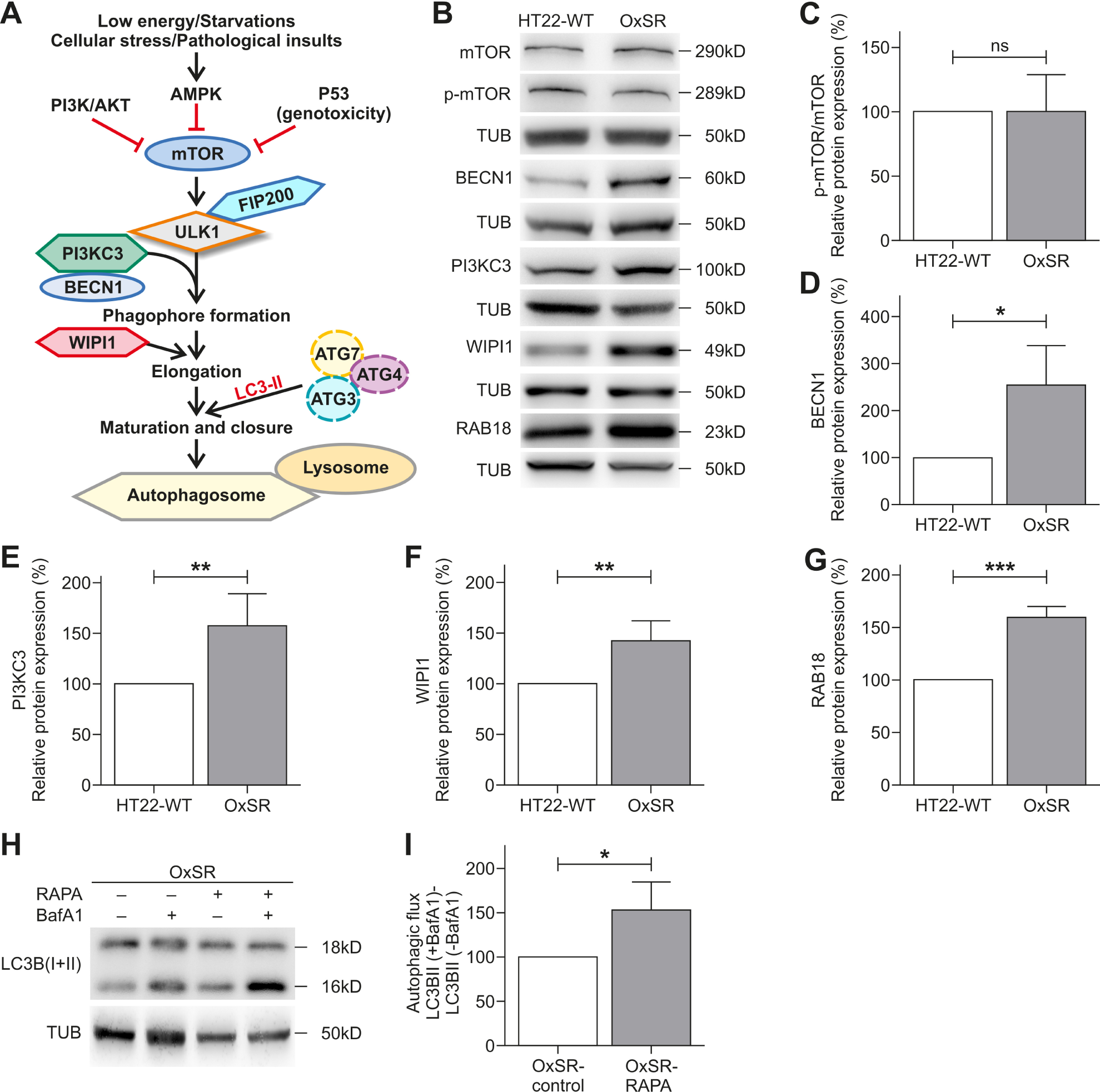
Adaptation to oxidative stress enhances expression of Beclin1, WIPI1, PI3KC3 and RAB18. (A) Scheme displays major regulators of canonical autophagy. (B) Protein extracts from untreated HT22-WT and OxSR cells were used to perform Western blot analysis for detection of indicated proteins. (C, D, E, F, G) The diagrams display the indicated protein levels after normalization to corresponding tubulin. Values represent mean ± S.E.M., n=3 to 5, *p<0.05, **p<0.01, *** p<0.001 and ns. for no significant statistical difference. Control HT22-WT cells were set to 100%. (H, I) OxSR cells were incubated simultaneously with 10 µM Rapamycin (RAPA) and 8 µM of BafA1 for 4 h and expression of the indicated proteins was detected by Western blotting. The autophagic flux was measured after densitometry analysis with LC3B-II levels normalized to tubulin. Values represent mean ± S.E.M., n=4, *p<0.05 and autophagic flux of untreated OxSR cells were set as control 100%.

Another upstream regulator of autophagy is the PI3KC3/BECN1 complex required for the recruitment of PI(3)P during autophagosome generation. Interestingly, we found an almost 2.5-fold higher BECN1 protein level in OxSR cells indicating that the BECN1 complex is highly active in oxidative stress-adapted cells (Figure 2B and D). WIPI1 acts downstream as PI(3)P effectors for proper membrane rearrangement [44], [45] and consistent with the observed increased protein level of BECN1, Western blot analysis indeed revealed also upregulated levels of both PI3KC3 and WIPI-1 in OxSR cells (Figure 2B, E and F). Moreover, RAB18 was found to be upregulated in its expression approximately 1.6-fold in OxSR cells (Figure 2B and G). The small Rab GTPase RAB18 is a known key regulator of ER-Golgi membrane trafficking, involved in cellular lipid metabolism and as recently shown also shown a positive modulator of autophagy [46], [47], [48]. Taken together, in OxSR cells we found a significant upregulation of the expression of distinct key proteins of canonical autophagy regulation comprising BECN1, WIPI1 and PI3KC3 as well as the positive autophagy regulator RAB18.

### 3.3 Autophagy qPCR array reveals a high upregulation of ATG9 mRNA in OxSR cells

Employing a qPCR array representing a wide range of genes involved in the direct regulation of various stages of the autophagic process itself as well as candidates that are not part of the core canonical autophagy machinery, we found a highly differential mRNA expression pattern comparing OxSR and WT cells. OxSR cells showed significantly higher mRNA expression of certain ATG proteins, including ATG10, ATG12, ATG5, as well as GABRAPL2 and AKT1S (Table 1). Moreover, consistently with our Western blot findings (see Figure 2), the qPCR also showed an elevated level of BECN1 and WIPI1 mRNA in OxSR cells (Table 1). Further, the expression of the lysosomal marker LAMP1 mRNA was upregulated in OxSR cells, confirming earlier observation in H2O2-resistant HT22 cells (Table 1) [28].

**Table 1:** Autophagy associated gene expression profiling of stress-resistant OxSR cells. Total RNA from HT22-WT and OxSR cells was characterized by using the mouse Autophagy Primer Library 1 (MATPL-1) and relative fold change in gene expression profile was analyzed by comparing with HT22-WT cells as control. Green numbers indicate an upregulation greater than 1.5-fold, red numbers indicate a downregulation lower than 0.6-fold. Values represent mean ± S.E.M., n=3, *p<0.05, **p<0.01, *** p<0.001 and NS for no significant statistical difference.

| HT22-WT vs. OxSR |  |  |  |
| --- | --- | --- | --- |
|  | Gene Name | Fold Change | Significant level |
| ATG1/ULK complex | Ulk2 | 0.4 | * |
|  | Ambra1 | 2.4 | NS |
| Other ATG | Atg10 | 2.3 | *** |
|  | Atg12 | 1.6 | *** |
|  | Atg16L2 | 0.6 | * |
|  | Atg4A | 3.3 | *** |
|  | Atg4B | 1.5 | NS |
|  | Atg4C | 2.6 | NS |
|  | Atg4D | 2.6 | NS |
|  | Atg5 | 2.3 | **** |
|  | Atg7 | 2.0 | NS |
|  | Atg9A | 1.9 | NS |
|  | Atg9B | 36.6 | * |
|  | Wipi1 | 1.7 | *** |
|  | Gabarapl1 | 1.9 | NS |
|  | Gabarapl2 | 1.6 | ** |
|  | Map1LC3A | 0.2 | *** |
| BECN1/PI3KC3 complex | Bcl2L1 | 5.4 | NS |
|  | Becn1 | 2.1 | ** |
|  | PIK3c3 | 1.7 | NS |
| Autophagy adaptor | Sqstm1 | 3.3 | *** |
| mTOR complex | Raptor | 2.2 | NS |
|  | Rictor | 1.8 | NS |
|  | AKT1s1 | 1.5 | ** |
| Lysosome associated | Dram | 1.3 | NS |
|  | Lamp1 | 3.4 | ** |
|  | Lamp3 | 2.3 | NS |
| others | Bire5 | 1.5 | * |
|  | Bnip3 | 6.1 | *** |
|  | Ddit3 | 2.1 | ** |
|  | Eif4ebp1 | 2.1 | *** |
|  | Eif4g1 | 1.9 | ** |
|  | Fkbp15 | 0.5 | NS |
|  | Gfi 1 b | 3.2 | NS |
|  | Gpsm1 | 1.9 | * |
|  | Letm1 | 3.3 | *** |
|  | Letm2 | 3.8 | *** |
|  | Rasd1 | 2.0 | NS |
|  | Sec16A | 2.3 | NS |
|  | Sec16B | 1.6 | NS |
|  | Sec23A | 2.0 | * |
|  | Sec23B | 2.2 | ** |
|  | Sh3glb1 | 2.0 | ** |
|  | Sh3glb2 | 3.4 | ** |
|  | Snx30 | 1.9 | NS |
|  | Tpr | 1.6 | * |
|  | Tpr53 | 0.5 | * |
|  | Ulk3 | 1.8 | NS |

Most significantly, we also found an over 30-fold upregulation of the mRNA expression of ATG9B in OxSR cells (Table 1). ATG9 proteins are essential for phagophore maturation (meaning early steps in autophagosome generation) and vesicle elongation and also determines the size of autophagosomes by acting as a lipid carrier delivering lipids to the growing phagophore [49], [50], [51], [52]. In addition, also a higher expression of the SEC23a and SEC23b mRNA was observed in OxSR cells (Table 1). SEC protein family members are important for the formation of specialized transport vesicles from ER to autophagosome biogenesis and selection of specific cargo molecules [53], [54].

The PCR mRNA array further revealed a significant increase in the mRNA expression of BNIP3 (6-fold) and BNIP3 is known to regulate mitophagy, the autophagic clearance process of damaged mitochondria (Table 1) [55]. LETM1 and 2, which are not directly related to autophagy but required to maintain mitochondrial ion homeostasis and morphology [56], [57] were also found to be highly upregulated in OxSR cells in their expression (Table 1) linking oxidative stress adaptation also to altered mitochondrial function; this link is further confirmed by full proteome analysis (see 3.7 below). Consistent with these data, the observed alteration of overall mitochondrial morphology in OxSR cells is also supported by highly elevated levels of DLP1, which is responsible for mitochondrial fission, as detected by Western blot analysis (Supplementary Figure S1C and D).

### 3.4 OxSR cells can handle more effectively proteotoxicity

Next, we investigated how effectively OxSR cells can handle disturbances of the proteostasis network. To reproducibly induce protein stress, we employed the compound canavanine (CV), an arginine analogue which disturbs protein homeostasis by interfering with protein translation. Western blot analysis revealed an increased accumulation of polyubiquitinated degradation-prone proteins in HT22-WT but not in OxSR cells after 12 h treatment of the cells with CV (Figure 3A and B). Since the proteasomal activity in OxSR cells was lower compared to HT22 WT cells (see below and Supplementary Figure S1E) the increased ability of the OxSR cells to cope with proteotoxic stress as indicated by the accumulation of polyubiquitinated proteins could be a direct result of an increased autophagic clearance activity.

**Figure 3:**
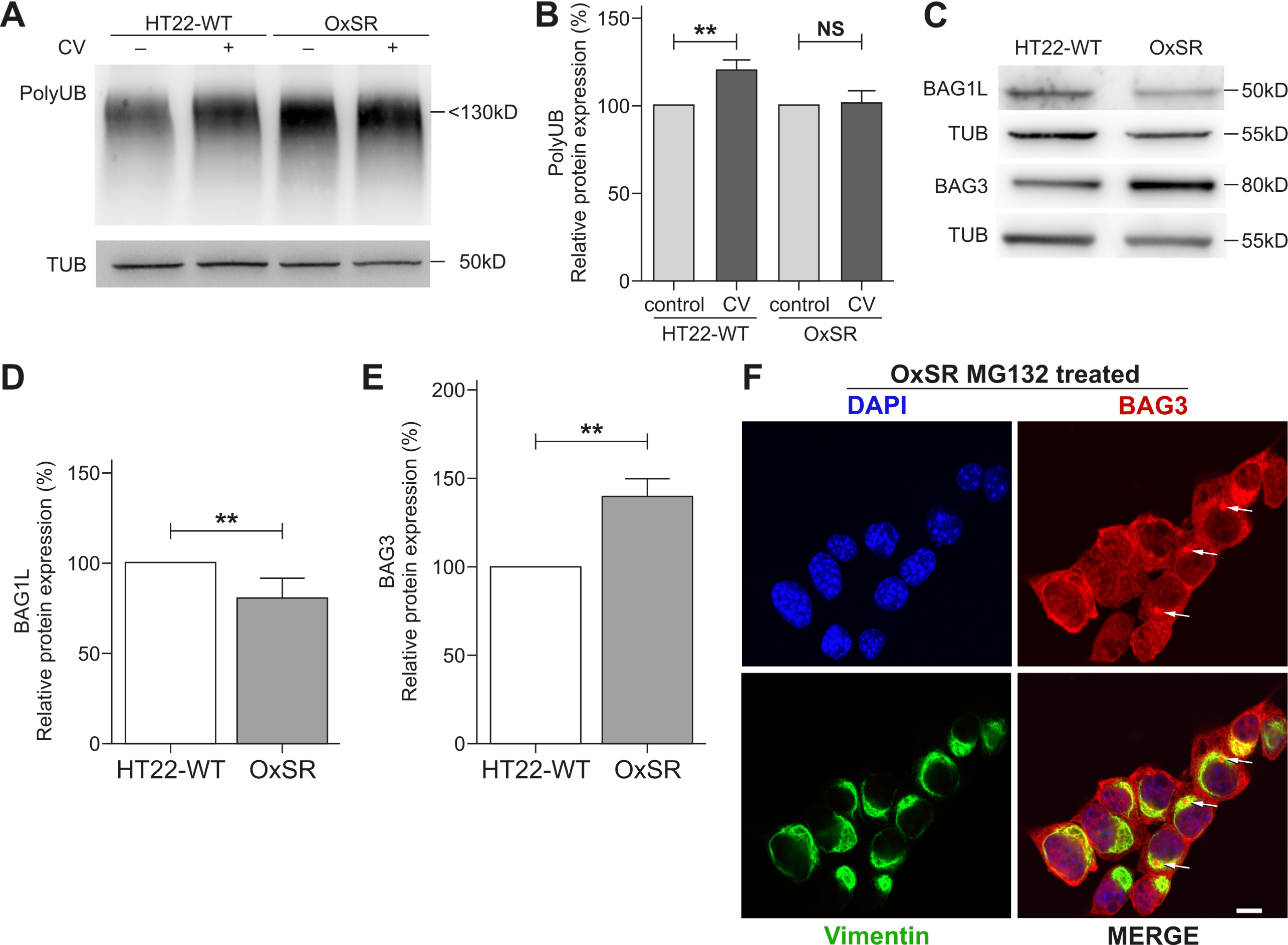
OxSR cells overcome proteotoxic stress more effectively than HT22-WT cells and show an altered BAG1/BAG3 expression. (A, B) protein extracts from untreated and canavanine- (CV-) treated samples of HT22-WT and OxSR cells were analyzed by Western blot for indicated proteins. Bar graphs represent changes in polyubiquitinated protein levels between untreated control and treated samples obtained by densitometric analysis after normalized to tubulin. Values represent mean ± S.E.M., n=3, **p<0.01 and NS for no significant statistical difference. Polyubiquitinated proteins in untreated cells were set as control 100%. (C) Protein extracts from untreated HT22-WT and OxSR cells were subjected to Western blot analysis for BAG1 and BAG3 expression; tubulin was used as loading control. (D, E) Quantification of the expression of BAG1L and BAG3 by densitometry analysis and normalization to tubulin. Values represent mean ± S.E.M., n=3 to 4, **p<0.01 and control HT22 cells were set to 100%. (F) OxSR cells were treated with MG132 and immunocytochemically analyzed for BAG3 positive aggresomes surrounded with cage like vimentin structure by confocal microscope. Scale bar: 10 µm.

### 3.5 BAG1/BAG3 expression switch in OxSR cells

In OxSR cells chronically adapted to oxidative stress we observed a switch from BAG1 to BAG3 expression as previously already seen in human primary fibroblasts and neurons under acute redox stress and during aging [21], [58]. BAG3 expression is increased in OxSR cells compared to HT22-WT cells, while the expression of BAG1L in OxSR cells was decreased as detected by Western blotting (Figure 3C-E); BAG1L is the same isoform of BAG1 which was detected to be decreased in aged cells or cells after acute oxidative stress treatment [21]. Taken together, BAG3 and BAG1 (isoform BAG1L) expression are reciprocally regulated in OxSR cells. Next, the proteasomal chymotrypsin-like activity in OxSR and HT22-WT cells was analyzed and found to be decreased in OxSR cells (Supplementary Figure S1E). These results suggest that OxSR cells depend more on the autophagic system (driven by BAG3 [21]) rather than on the proteasomal system (driven by BAG1) for protein degradation. To further confirm the recruitment of BAG3 into the proteostasis network of OxSR cells proteasomal activit was inhibited by MG132, an efficient stimulus of protein aggresome formation [21]. The observed colocalization of BAG3 with vimentin-positive perinuclear aggresomes after pharmacological proteasome inhibition in the OxSR cells (Figure 3F) suggests an enhanced activity of BAG3-mediated aggresome targeting and autophagy.

### 3.6 BAG3 overexpression in HT22-WT cells leads to increased autophagic flux and lysosomal rearrangement, BAG3 depletion in OxSR cells results in reduced autophagic flux

To clarify a direct participation of BAG3-mediated selective macroautophagy in the enhanced overall autophagic activity and the lysosomal phenotype of OxSR cells, we overexpressed BAG3 in HT22-WT cells. HT22-WT cells following transfection with a BAG3 expression construct showed a significantly increased autophagic flux (Figure 4A, B and D). Intriguingly, along with overexpression of BAG3 also the protein levels of the lysosomal membrane protein LAMP2 were found to be strongly enhanced (Figure 4A and 4C). This suggests that BAG3 expression can have a direct effect on lysosomes. Finally, lysosomal compartments were analyzed by LysoTracker staining in HT22-WT (as controls), HT22-WT cells transfected with empty vector (HT22-WTev), HT22-WT cells overexpressing BAG3 (HT22-WTBAG3) and OxSR cells. Perinuclear lysosomes were found in HT22-WT and HT22-WTeV cells (controls), however, BAG3-overexpressing HT22-WTBAG3 cells showed an elevated number of lysosomes with cellular distribution comparable to that in OxSR cells (Figure 4I). Quantification showed that also the total number of cells showing distinguishable acidic vacuolar structure is significantly elevated in HT22-WTBAG3 compared to HT22-WT and HT22-WTeV and is comparable to OxSR cells (Figure 4H and I).

**Figure 4:**
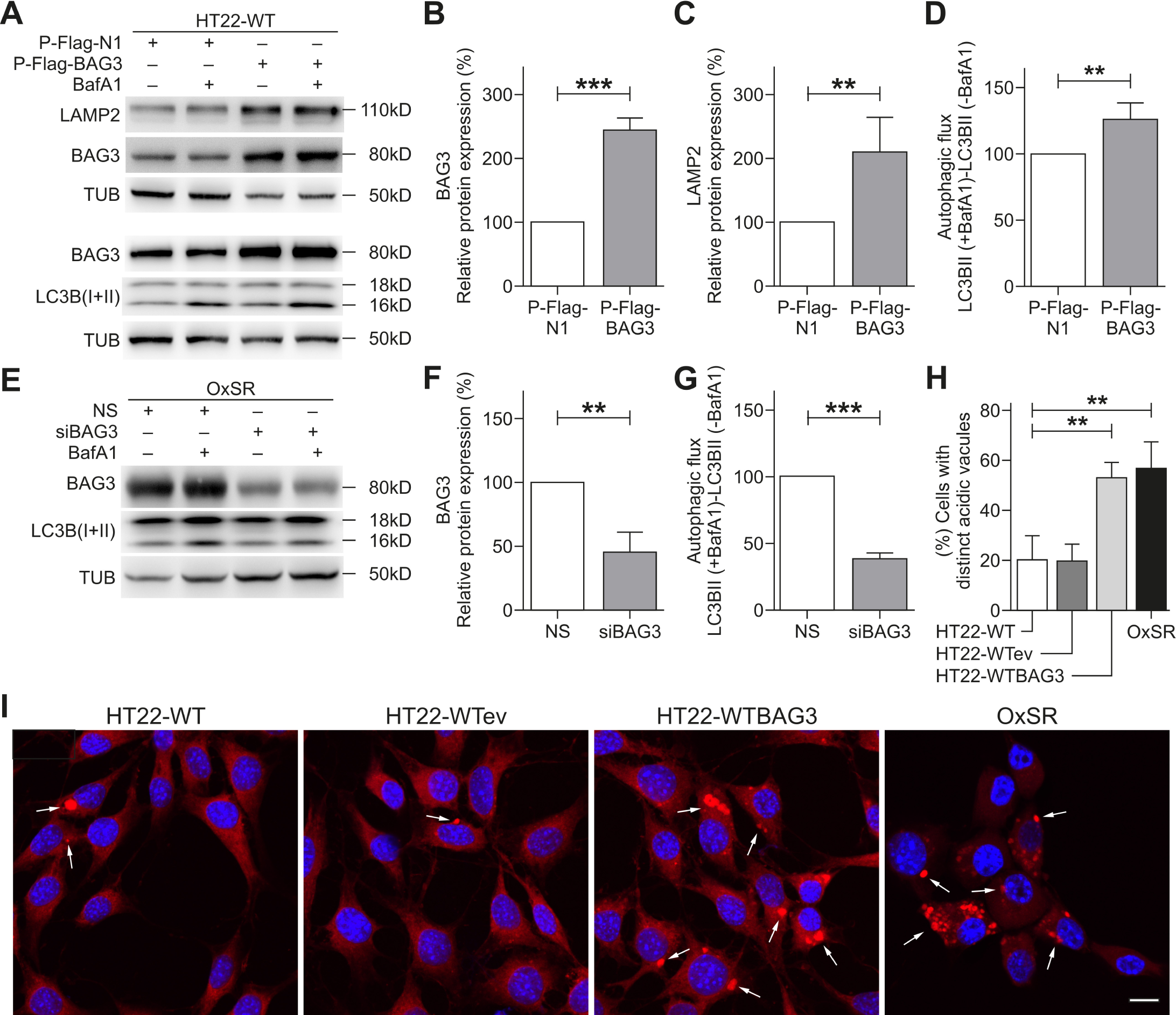
BAG3 is required for autophagic activity in OxSR lines and BAG3 overexpression promotes autophagy in HT22-WT cells. (A, B, C, D) HT22-WT cells transfected for 48 h with empty vector (P-Flag-N1) or BAG3 overexpression vector (P-Flag-BAG3) and treated with vehicle or with BafA1 for 6 hr before the cell lysis. Protein extracts were analyzed via Western blotting for indicated proteins. Quantification of LAMP2, BAG3 expression and autophagic flux were prepared by densitometric analysis after normalization of protein levels to tubulin. Values represent mean ± S.E.M., n=4, **p<0.01 and *** p<0.001. Expression in empty vector transfected cells were set to 100%. (E, F, G) OxSR cells were transfected with nonsense siRNA (NS) and BAG3 siRNA (siBAG3) for 48 h. Protein extracts from vehicle or BafA1-treated cells were subjected to Western blot analysis. Quantification of BAG3 expression and autophagic flux were prepared after densitometric analysis and normalization to tubulin. Values represent mean ± S.E.M., n=3, **p<0.01 and *** p<0.001. Expression in nonsense siRNA transfected cells were set to 100%. (H) Quantitative bar graph shows number of cells with distinct acidic vacuoles found with LysoTracker staining in untreated HT22-WT, HT22 empty vector transfected (HT22-WTeV), HT22 overexpressing BAG3 (HT22-WTBAG3) and OxSR cells. Values represent mean ± S.E.M., n=4 with 3 to 8 images of different microscopic fields was counted in each day for every experimental condition, **p<0.01. (I) Representative confocal images of LysoTracker red staining (LT) in untreated HT22-WT, HT22-WTeV, HT22-WTBAG3 and OxSR cells are shown indicating the changes in lysosomal phenotype. Arrows indicate cells containing distinct acidic vacuolar structure. DAPI (blue) was used to stain DNA. Scale bars: 10 µm.

In addition, after knockdown of BAG3 in OxSR cells, autophagic flux was analyzed by determining the accumulation of LC3B-II after treatment with BafA1 by Western blot. Upon knockdown of BAG3 via siRNA a significant reduction of the autophagic flux was found in OxSR cells (Figure 4E-G), whereas the downregulation of BAG3 by siRNA showed no change in autophagic flux of HT22-WT cells (data not shown). Taken together these results strongly support the role of BAG3 and BAG3-mediated selective macroautophagy in the adaptation of OxSR cells to chronic redox stress.

### 3.7 Total proteome analysis underlines differential expression of proteins involved in autophagy, mitochondrial function and metabolism

Finally, to analyze changes in the global proteome in OxSR cells (as compared to HT22-WT) we performed a bottom-up mass spectrometry-based discovery proteomics analysis. Label-free quantification (LFQ) analysis of the triplicates of both cell lines employing a false discovery rate (FDR) of 1% identified a total of 1121 proteins (the complete dataset is included in Supplementary file S4A “Proteomics data”). Subsequently, we identified the proteins that exhibited significant variations between OxSR and HT22-WT cells. For this, the experimental reproducibility of the LFQ between the replicates was examined employing Pearson’s correlation coefficients, which demonstrated excellent biological reproducibility with an average R value of 0.97 ± 0.01 and R of 0.93 ± 0.01 between the two cell lines. Data is represented in the scatter plots of the intensity distribution in one replicate relative to another (Supplementary file S4B “Proteomics data”).

Among the identified proteins, 176 were differentially expressed, with 87 up-regulated and 89 down-regulated proteins (Supplementary file S4C “Proteomics data”). Further clustering of these differentially expressed proteins depicts the distinct patterns of the proteome presented as a heat map with unsupervised hierarchical clustering (Figure 5A). Closer examination of these protein clusters revealed significant changes in several cellular functions (Figure 5B and Supplementary file S4D “Proteomics data”). Endoplasmic reticulum stress response, fatty acid metabolism, synthesis of reactive oxygen species, glycolysis, mitochondrial disorder, oxidative stress, autophagic cell death, neuronal cell death, autophagy and cell survival of neurons represent the top ten most significant functional changes implicated in OxSR cells in comparison to the oxidative stress-sensitive HT22-WT cells.

**Figure 5:**
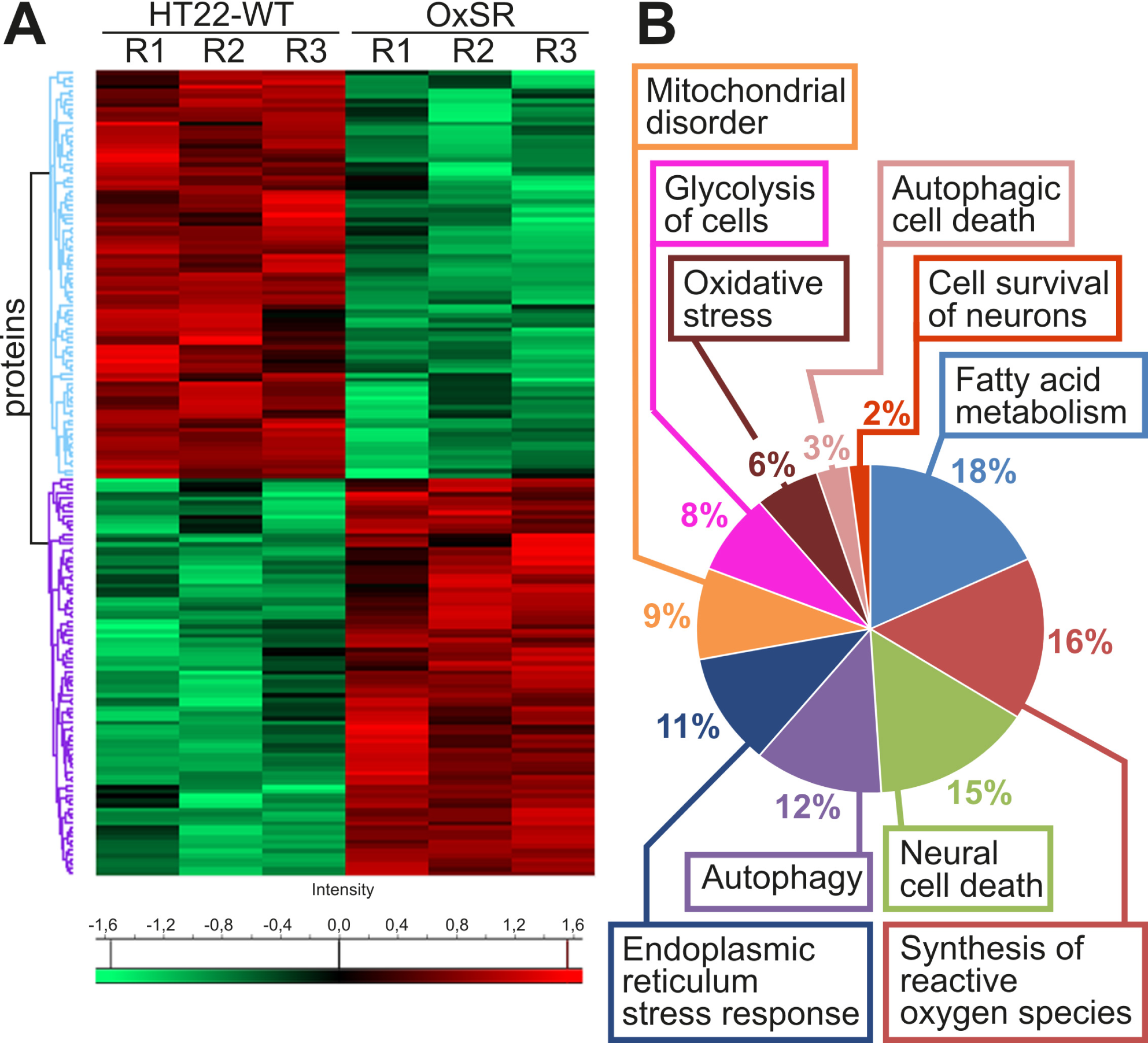
Comparative proteomics analyses of the HT22-WT and OxSR cells. (A) Hierarchical clustering of the 176 differentially expressed proteins depicted as a heat map distinguishes two major clusters of proteins. The first cluster represents significantly down-regulated (green) and the second cluster up-regulated (red) proteins in OxSR cells compared to HT22-WT. R1 to R3 represent the biological replicates. (B) Pie chart displays cellular functions and abundance of proteins which are differentially expressed in OxSR cells compared to HT22-WT cells. The detailed list of proteins in each cellular functional group is provided with Supplementary file S4D “Proteomics data” where arrows depict the up- and down-regulation of the proteins, respectively.

Finally, in an effort to further explore protein-protein interaction pathways (PPIs) of the proteins identified to be differentially expressed, functional pathway enrichment was determined employing the Ingenuity Pathway Analysis (v01-04, IPA; Ingenuity QIAGEN Redwood City, CA) tool. Consistently, a large majority of these proteins were found to be actively involved in the process of autophagy (14 proteins) but also linked to mitochondrial function and disorders (16 proteins) and oxidative stress/redox signaling (7 proteins) (Supplementary Figure S2 and the complete list to be found in Supplementary file S4E “Proteomics data”).

## 4. Discussion

A disturbed protein homeostasis, proteotoxic and oxidative stress are hallmarks of neurodegeneration and aging [59], [60], [61]. Autophagy is described as a physiological process involved in antioxidant defense [62] and autophagy deficiency clearly results in neuronal loss and neurodegeneration in mice [63]. On the other hand, induction of autophagy is shown to alleviate oxidative stress in myocardial ischemia/reperfusion Injury [64].

The adaptation to oxidative stress is a highly relevant process and some adaptive pathways focusing on antioxidant defense systems and mitochondrial and metabolic adaptations have been described to be constantly activated in neuronal cells resistant to oxidative stress as well as in AD tissue [26], [29], [6]. Moreover, oxidative stress resistance has been linked to failure of anticancer therapy and here a particular role of upregulated autophagy and BAG3 has been proposed in various tumor types [16], [65], [66], [67].

Based on the knowledge that dysregulated autophagy is actively involved in neurodegeneration [68], [69] combined with previous findings on mouse hippocampal HT22 cells fully adapted to high concentrations of H2O2 (OxSR cells) [28], we describe here in detail the molecular changes associated with redox stress adaptation in neuronal cells. To do so, we engaged different expression and functional studies including morphological parameters obtained by TEM, investigation of autophagic flux and global proteome analysis using mass spectrometry.

We found a significantly increased autophagic activity in the cells adapted to chronic oxidative stress (OxSR cells) (Figure 1) and on the other hand a reduced proteasomal function (Supplementary Figure S1E) suggesting that autophagy may serve as the preferred protein degradation pathway during oxidative stress challenge. We identified several key proteins involved in autophagy regulation that showed a significantly altered expression. ROS production i.e. oxidative stress is known to inhibit mTORC1 [64], [70], the major regulatory complex for autophagy induction. Interestingly, in OxSR cells the increase in autophagy under chronic oxidative stress is not mediated via mTOR inhibition, whereas the classical BECN1/PI3K3C3 complex obviously is involved and highly upregulated in their expression in OxSR cells (Figure 2). Moreover, RAB18 expression is enhanced in OxSR cells (Figure 2). This finding is consistent with the recent description of RAB18 being a positive autophagy regulator [48] and a role of RAB18 in lipid metabolism, which enables enhanced lipid recruitment for autophagosome formation [71].

In qPCR analysis we observed a massively increased expression of ATG9B in OxSR cells (Table 1). ATG9 is an essential ATG family protein required for phagophore formation and exists in two homologues referred to as ATG9A and ATG9B. Trafficking of ATG9 from the Golgi apparatus to the site of autophagosome formation following autophagy induction (for instance by starvation) is reported to be dependent on PI3KC3 activity and negatively affected by PI3KC3 inhibition [72]. Therefore, it could be argued that upregulated ATG9B may work in concert with PI3KC3 in OxSR to promote autophagy. Leow et al. recently suggested that also sublethal acute oxidative stress can specifically induce ATG9B expression [73]. Furthermore, it is reported that ATG9 not only just acts as an autophagic membrane protein, but also upregulates autophagy induction via the JNK pathway by interacting with tumor necrosis factor receptor-associated factor 2 (dTRAF2) under oxidative stress condition [74]. In fact, ATG9 could be a key factor for the observed significantly enhanced autophagy activity in OxSR cells, since it can ensure the supply with membrane fragments early in autophagosome formation [75].

During cellular stress, the HSP70 chaperone system is a key player to maintain protein homeostasis together with its co-chaperones BAG1 (for proteosomal degradation) and BAG3 (for autophagic degradation). OxSR cells demonstrate an upregulated BAG3 accompanied with downregulated BAG1L expression (“BAG1/BAG3-switch”) (Figure 3) and cope with a disturbed proteostasis induced by canavanine more effectively. Also BAG3-positive aggresomes are found upon proteasomal inhibition as previously observed (Figure 3) [23]. Most interestingly, the overexpression of BAG3 causes a lysosomal rearrangement in HT22-WT cells that closely resembles the distribution (and numbers) of lysosomes seen in OxSR cells (Figure 4). The observed rearrangement of the lysosomal compartment by BAG3 may then have also functional consequences on the autophagic flux since LC3B-II flux was increased in HT22-WT cells following overexpression of BAG3 (Figure 4). Our results are consistent with recent observations that the lysosomal biogenesis is increased under sub-lethal acute oxidative stress and proposed to be mediated via a selective chaperone-mediated pathway [73]. Furthermore, the BAG3 siRNA knockdown significantly hampers the autophagic activity exclusively in OxSR cells and therefore, identifies BAG3 as one key positive autophagy modulator in OxSR cells (Figure 4). These data underline a strong role of BAG3 in boosting autophagy in OxSR cells but shows also for the first time that the expression switch from BAG1 to BAG3 and the functional switch from proteasomal pathway to BAG3-mediated macroautophagy can be of permanent nature under stable adaptation to oxidative stress as observed before during acute redox stress and aging [21], [58]. It is important to note here that neither knock-down of BAG3 via siRNA nor a general autophagy inhibition with bafilomycin A1 reversed the resistance phenotype (data not shown) strongly arguing that during long-term adaptation to redox stress actually a combination of different pathways may ensure oxidative stress resistance. This is strongly supported by our full proteome analysis as well as by previously identified factors that partially contribute to oxidative stress resistance such as NF-κB, sphingolipids and increased levels of antioxidant enzymes [26], [27], [28]. Indeed, employing MS-based global proteome analysis comparing OxSR and HT22-WT cells several proteins were identified to be differentially expressed and significant changes in several key pathways in addition to autophagy were revealed e.g. alteration in the ER stress response, fatty acid metabolism, glycolysis, oxidative stress response and survival of neurons, mitochondrial pathways and disorders. This argues towards the involvement of a wide array of changes involved in redox stress adaptation (Figure 5) and demonstrates also the plasticity and adaptability of various cellular pathways during chronic redox stress.

In summary we propose that upregulated autophagy in OxSR cells follows an mTOR-independent pathway. BECN1/PI3KC3 and ATG9B, critical proteins for autophagy induction and downstream execution respectively are highly upregulated during adaptation to oxidative stress. Moreover, the HSP70-cochaperone BAG3 is not only increased in its expression, but it is also active to participate in the upregulated selective macroautophagy in OxSR cells and also directly impacts on the lysosomal rearrangement. OxSR cells described here in detail may be a useful model to study these adaptive pathways and may also represent a tool to decipher further factors and interacting pathways able to boost autophagic flux in neuronal cells. Altogether analyzing the process of adaptation to chronic redox stress reveals potential novel therapeutic targets mediating a robust adaptability to long-term oxidative stress conditions and a disturbed proteostasis as important hallmarks of neurodegenerative disease and aging.

## Abbreviations

ATG: Autophagy related
BAG1: BCL2 Associated Athanogene 1
BAG3: BCL2 Associated Athanogene 3
BECN1: Beclin1
BNIP3: BCL2 interacting protein 3
BafA1: Bafilomycin A1
CTSD: Cathepsin D
CV: Canavanine
DLP1: Dynamin-like protein 1
Hsp70: Heat shock protein 70
LAMP1: Lysosomal-associated membrane protein 1
LAMP2: Lysosomal-associated membrane protein 2
LC3: Light chain 3 protein
LETM: Leucine zipper and EF-hand containing transmembrane protein
LFQ: Label-free quantification
mTOR: Mammalian target of rapamycin
MTT: (3-(4,5-Dimethylthiazol-2-yl)-2,5-Diphenyltetrazolium Bromide)
NS: Nonsense
PIK3C3: Class III PI3-kinase
PolyUB: Polyubiquitin
RAB18: Member RAS oncogene
Rapa: Rapamycin
siRNA: Small interfering RNA
TUB: Tubulin
TFEB: Transcription factor EB
TEM: Transmission electron microscopy
WIPI1: WD repeat domain phosphoinositide-interacting protein 1

## Acknowledgment

This study was supported by grants of the CRC1177 of the Deutsche Forschungsgemeinschaft (DFG) and the Peter Beate Heller Foundation to CB and Foundation Fighting Blindness (FFB) (PPA-0717-0719-RAD) (to UW). Caroline Manicam is supported by a grant from the DFG (MA 8006/1-1). The authors are grateful to Milena Rossmanith and Katharina Träger for their assistance with sample preparation for mass spectrometry analyses and to Andreas Kern, Jana Schepers and Albrecht Clement for critical reading.

---

**Supplementary figure S1:**
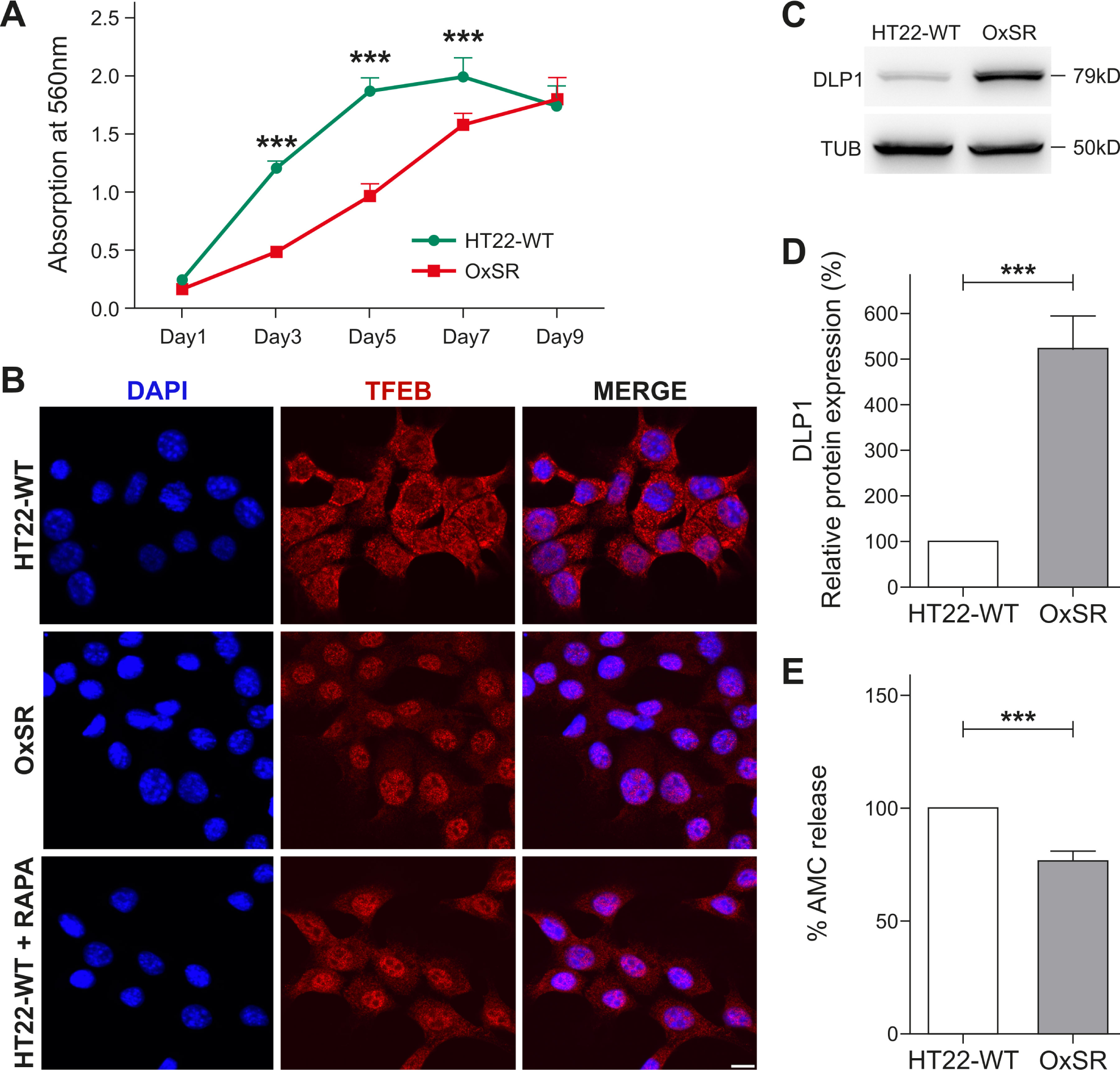
Results from cell proliferation assay, immunocytochemical analysis of TFEB, expression of mitochondrial protein DLP1 and Proteasomal activity. (A) Proliferation/growth rate of HT22-WT and OxSR cells were measured by MTT assay over nine alternate days. Values represent mean ± S.E.M., n=3, *** p<0.001 (B) Immunocytochemical analysis reveals a distinct nuclear translocation of TFEB in OxSR cells compared to cytosolic distribution in HT22-WT under basal condition. HT22-WT cells are treated with 10 µM Rapamycin for 2h to induce TFEB nuclear translocation as a positive control experiment (HT22-WT+RAPA). Scale bars: 10 μm. Images were acquired by confocal microscopy. (C) Protein extracts from untreated samples of HT22-WT and OxSR cells were analyzed by Western blotting for DLP1 expression. Tubulin was used as loading control. (D) Bar graph shows changes in DLP1 levels obtained by densitometry analysis and normalization to Tubulin. Values represent mean ± S.E.M., n=3, *** p<0.001. Control HT22-WT cells were set to 100%. (E) Proteasomal activity of HT22-WT and OxSR is measured using fluorescence signal by release of AMC from proteasomal substrate Suc-LLVY-AMC by proteasomal enzyme activity. Values represent mean ± S.E.M., n=5, *** p<0.001. Control HT22-WT cells were set to 100%.

**Supplementary figure S2:**
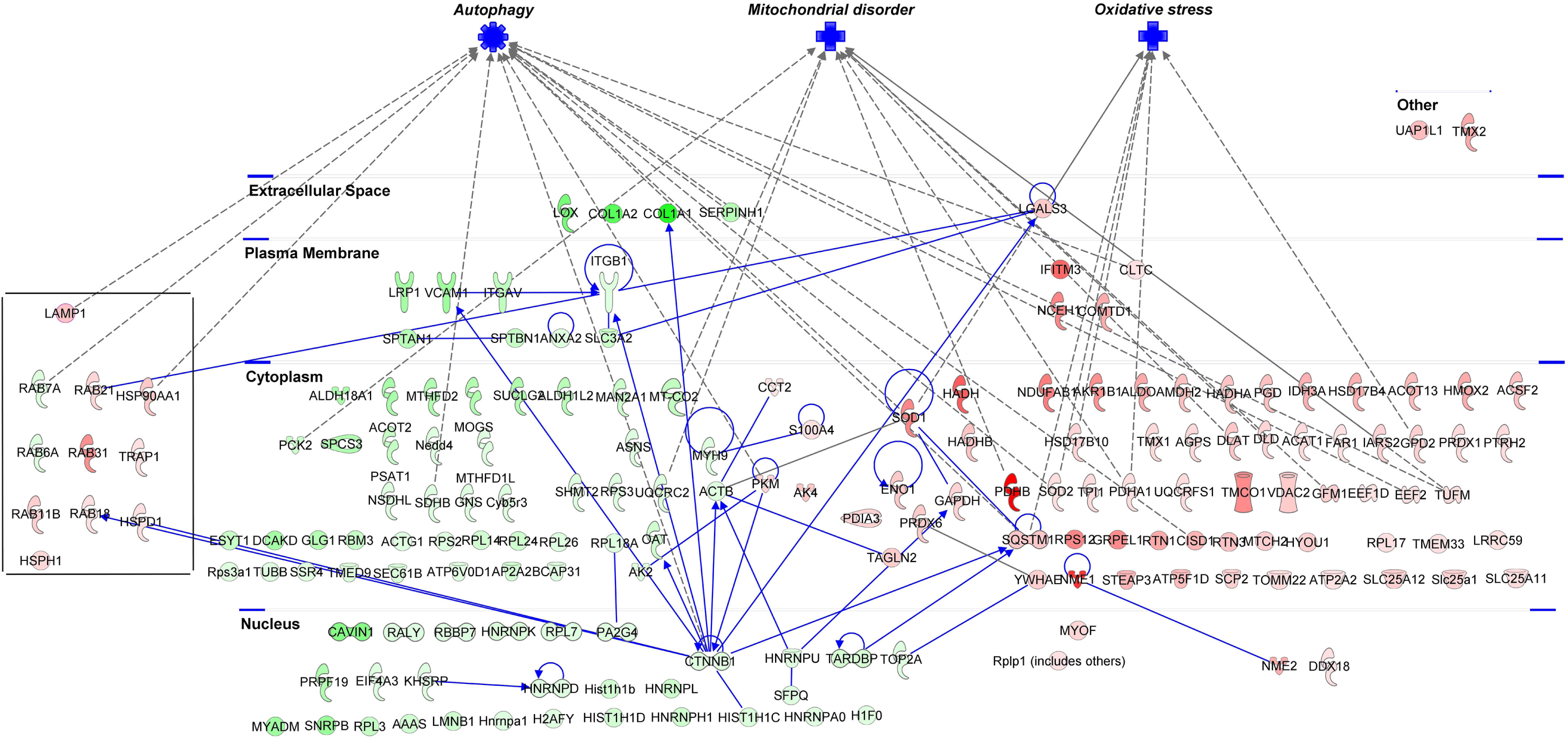
PPI Networks of the significantly differentially expressed proteins in OxSR cells. The protein-protein interaction (PPI) network of the differentially expressed proteins analyzed by IPA analysis demonstrated the involvement of three major protein clusters in autophagy, mitochondrial disorders and oxidative stress-related processes. Red and green colors of the molecules indicate up- and down-regulation of the proteins, respectively, and different intensities correspond to the magnitude of expression change of the individual proteins. The proteins were further distinguished according to their cellular localization and molecular functional classes with different shapes representing enzymes, ion channels, peptidases, transmembrane receptors etc. The detailed result is provided in Supplementary file S4E “Proteomics data”.

**Supplementary figure S3: Proteomics methods.**

Details of the materials, methods, statistics and software employed for proteomics analysis is described here.

**Supplementary figure S4:**
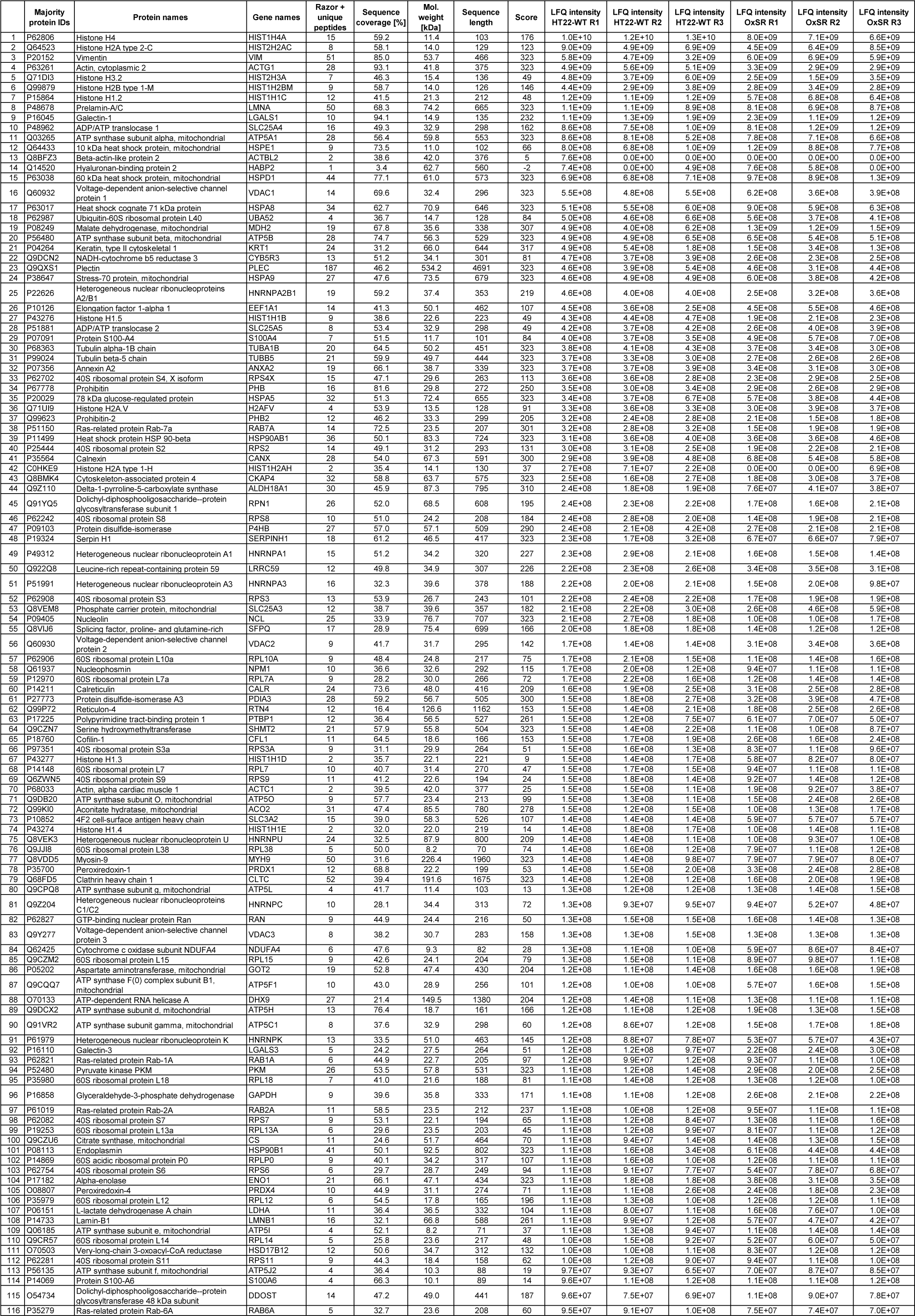
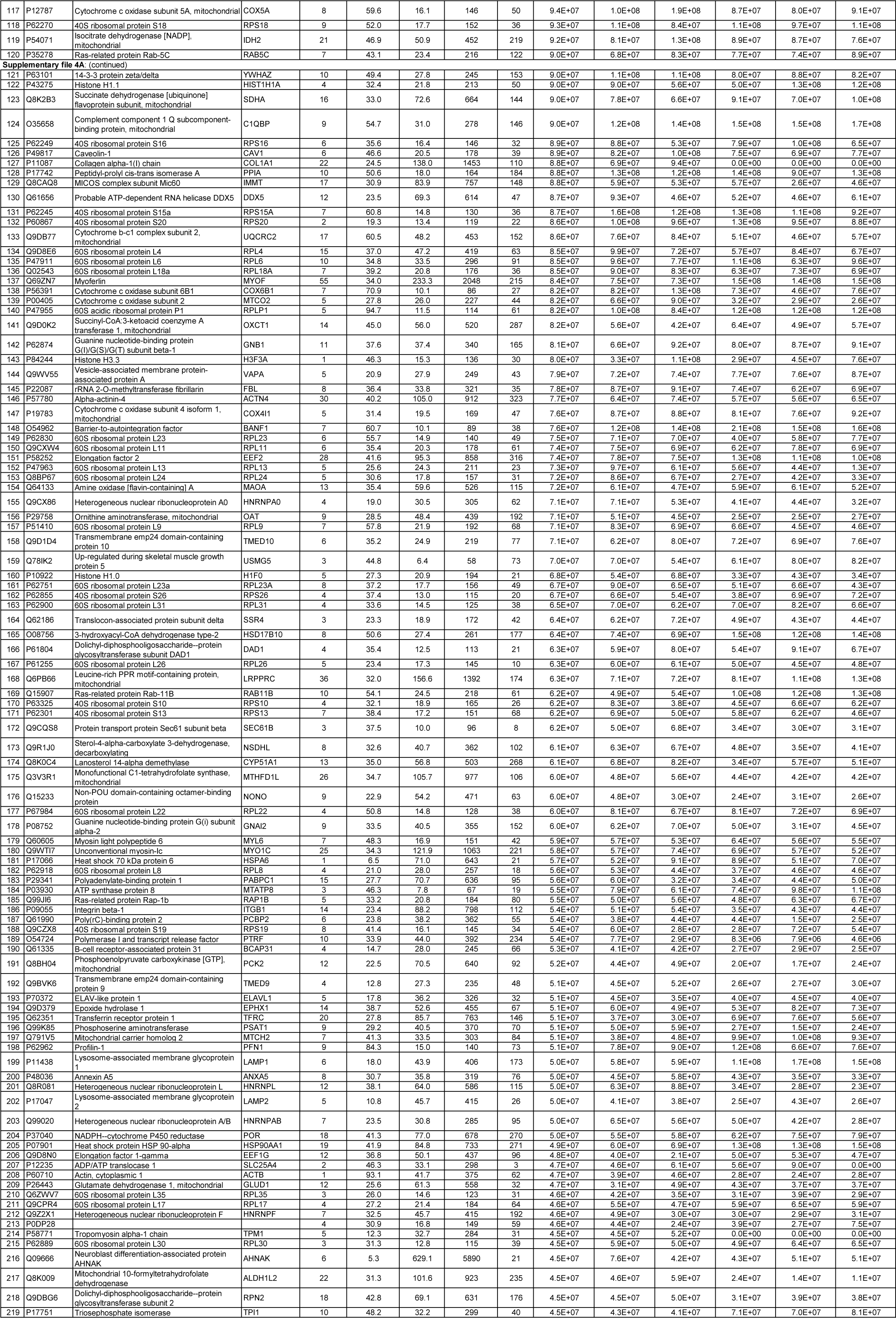
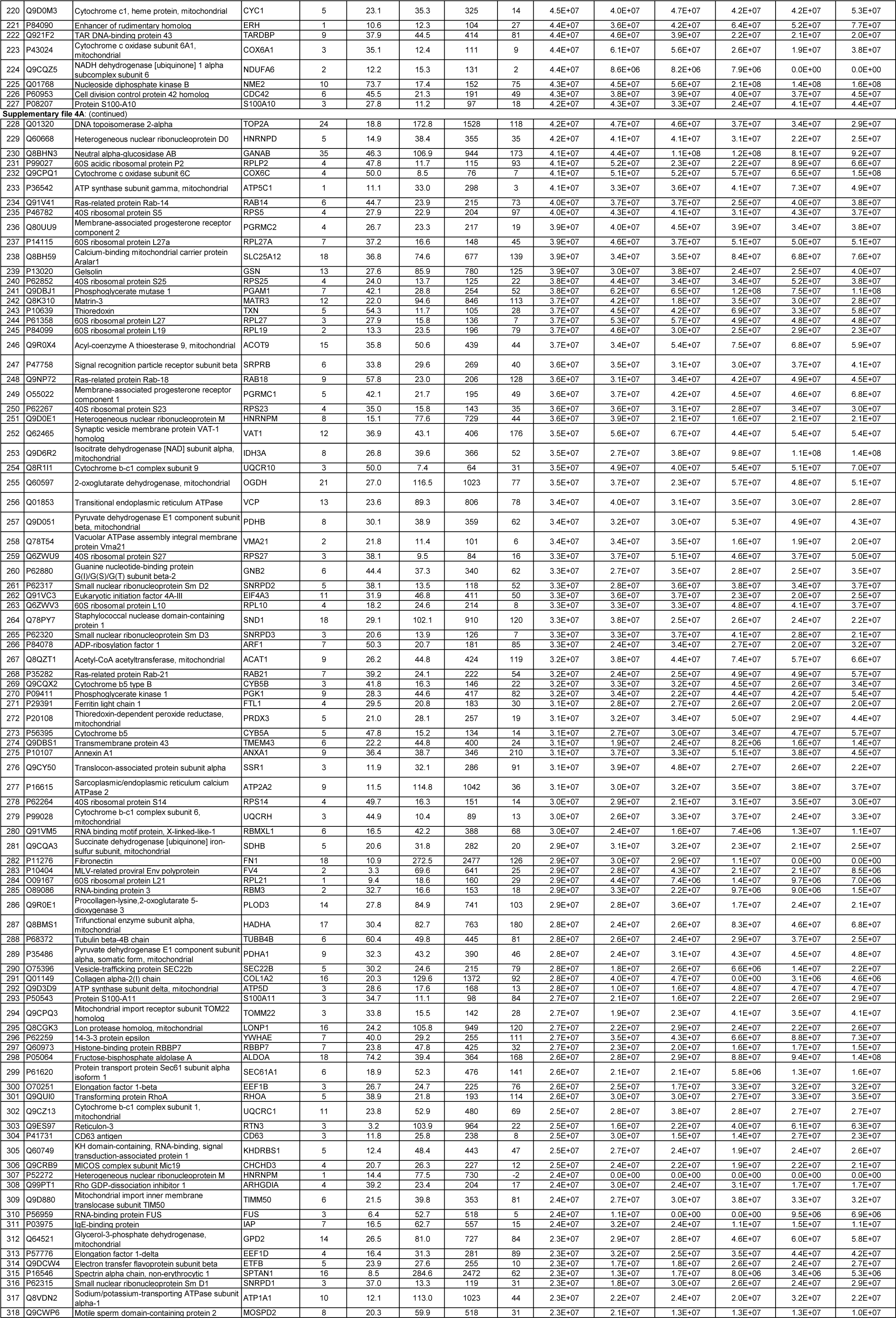
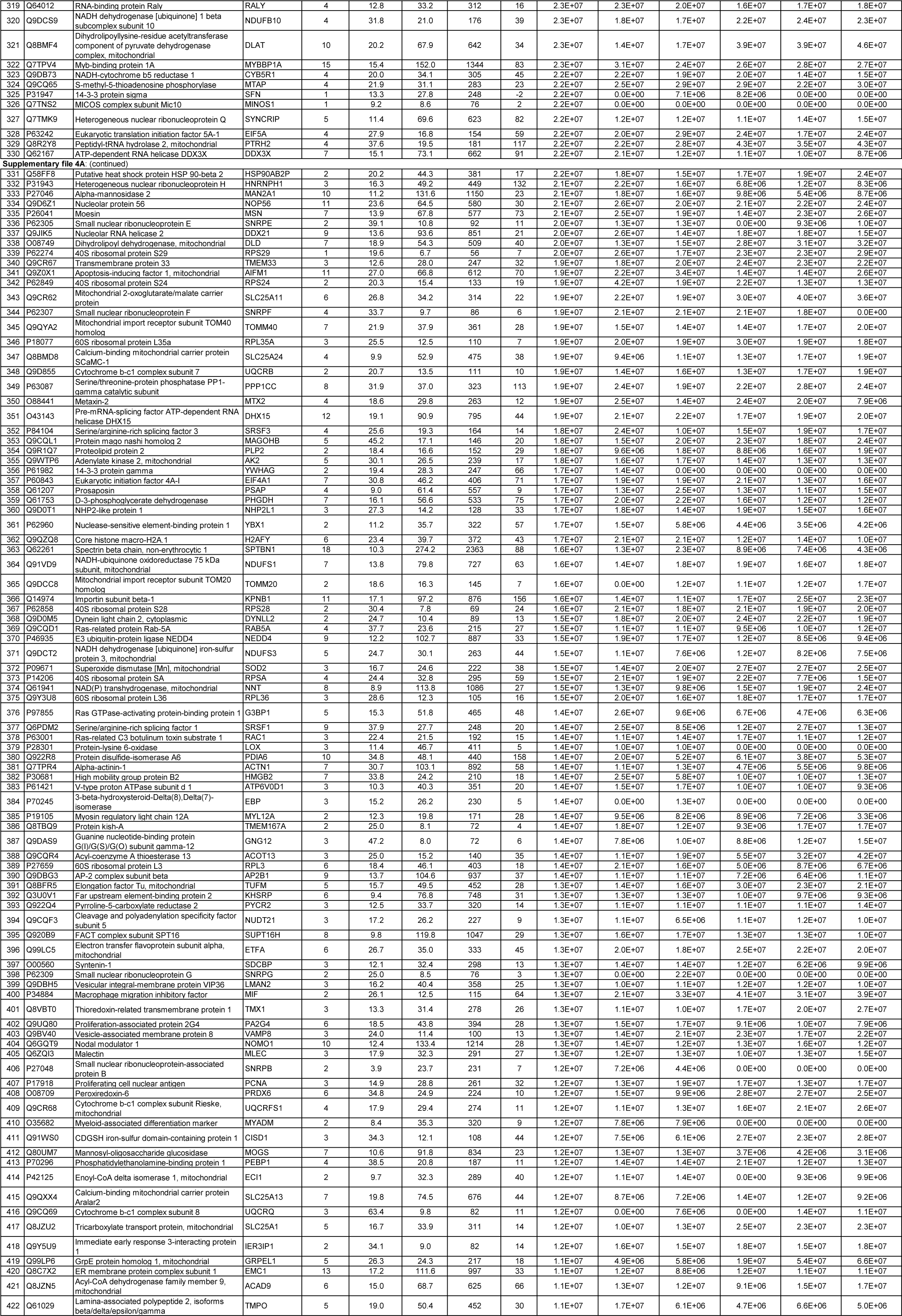
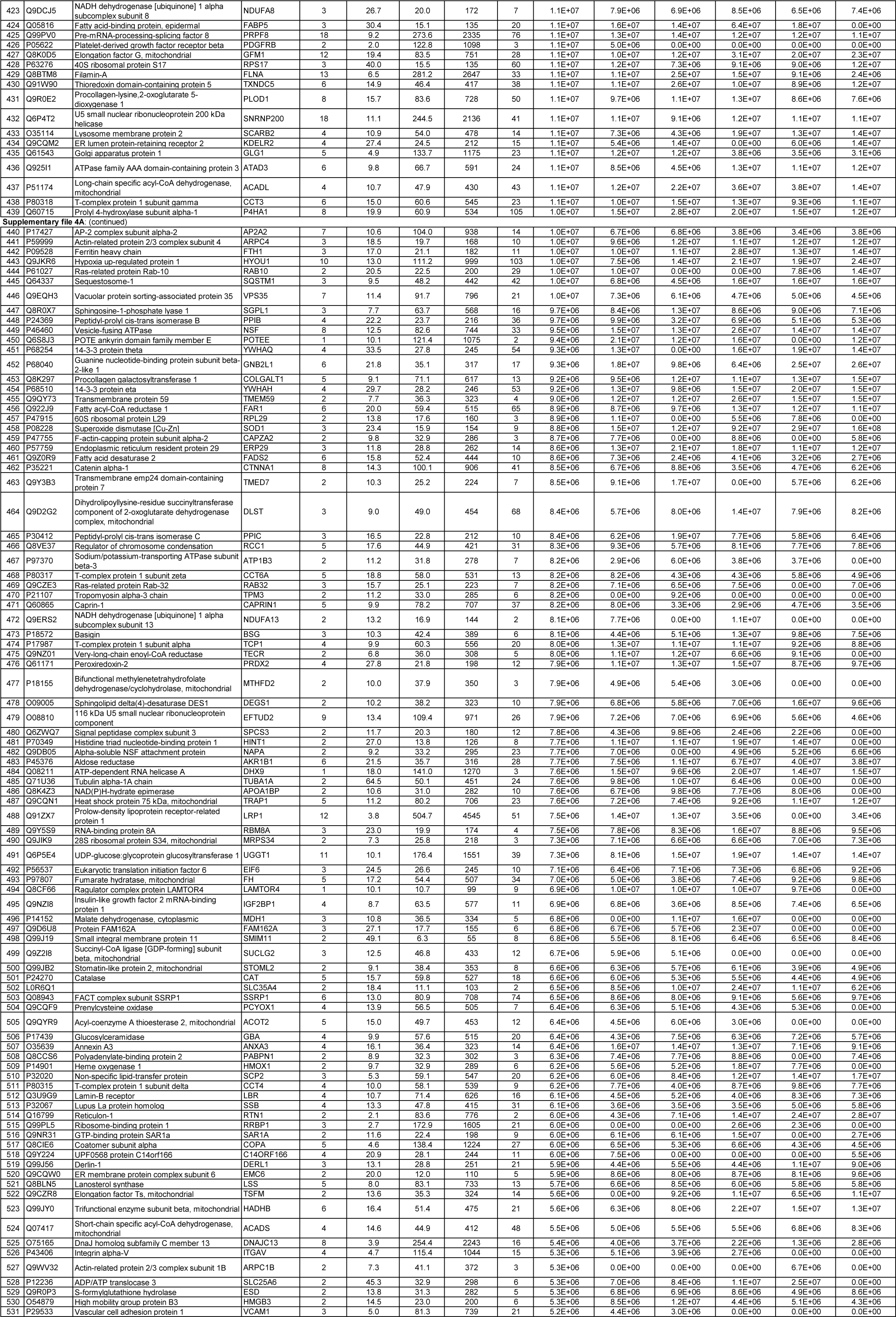
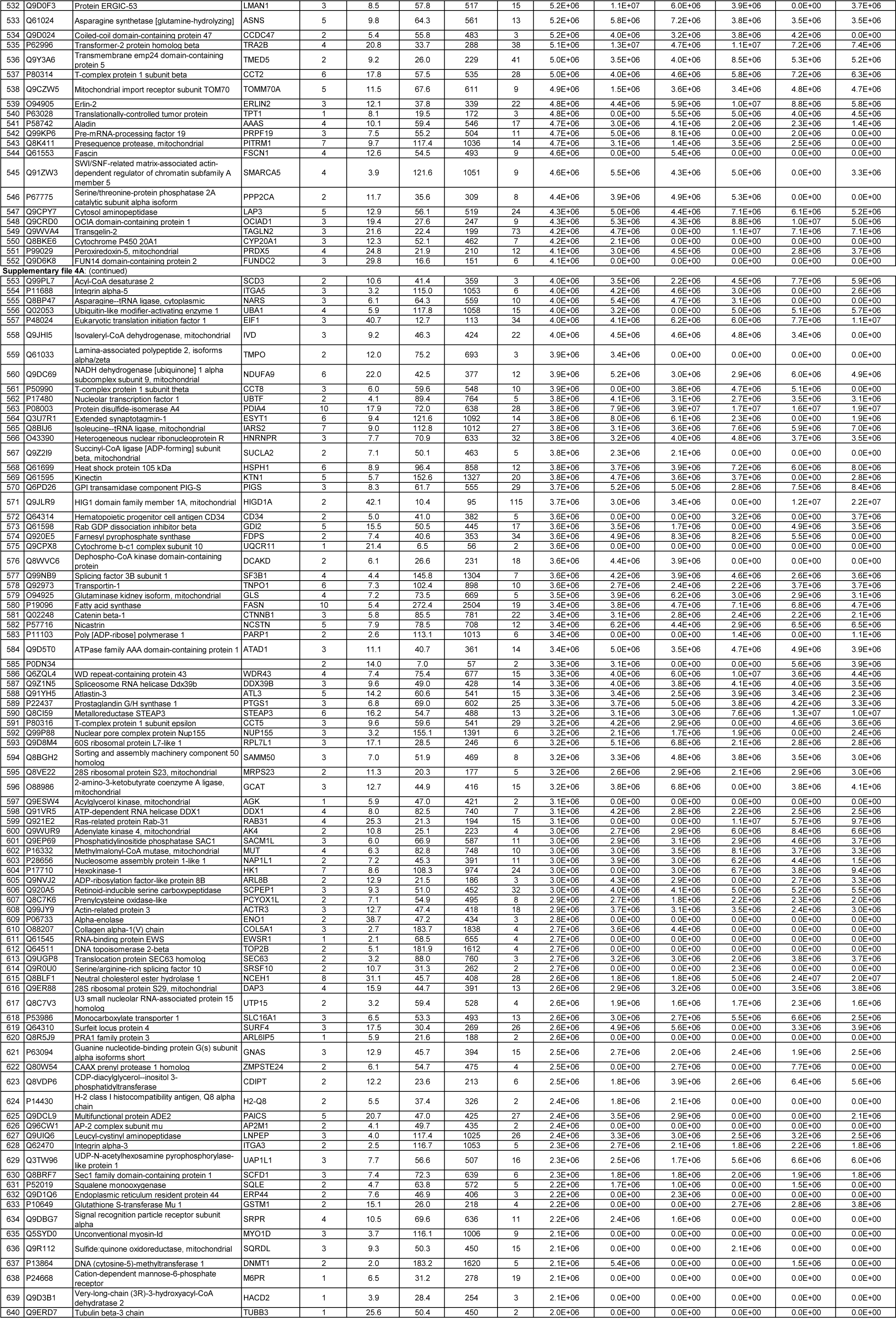
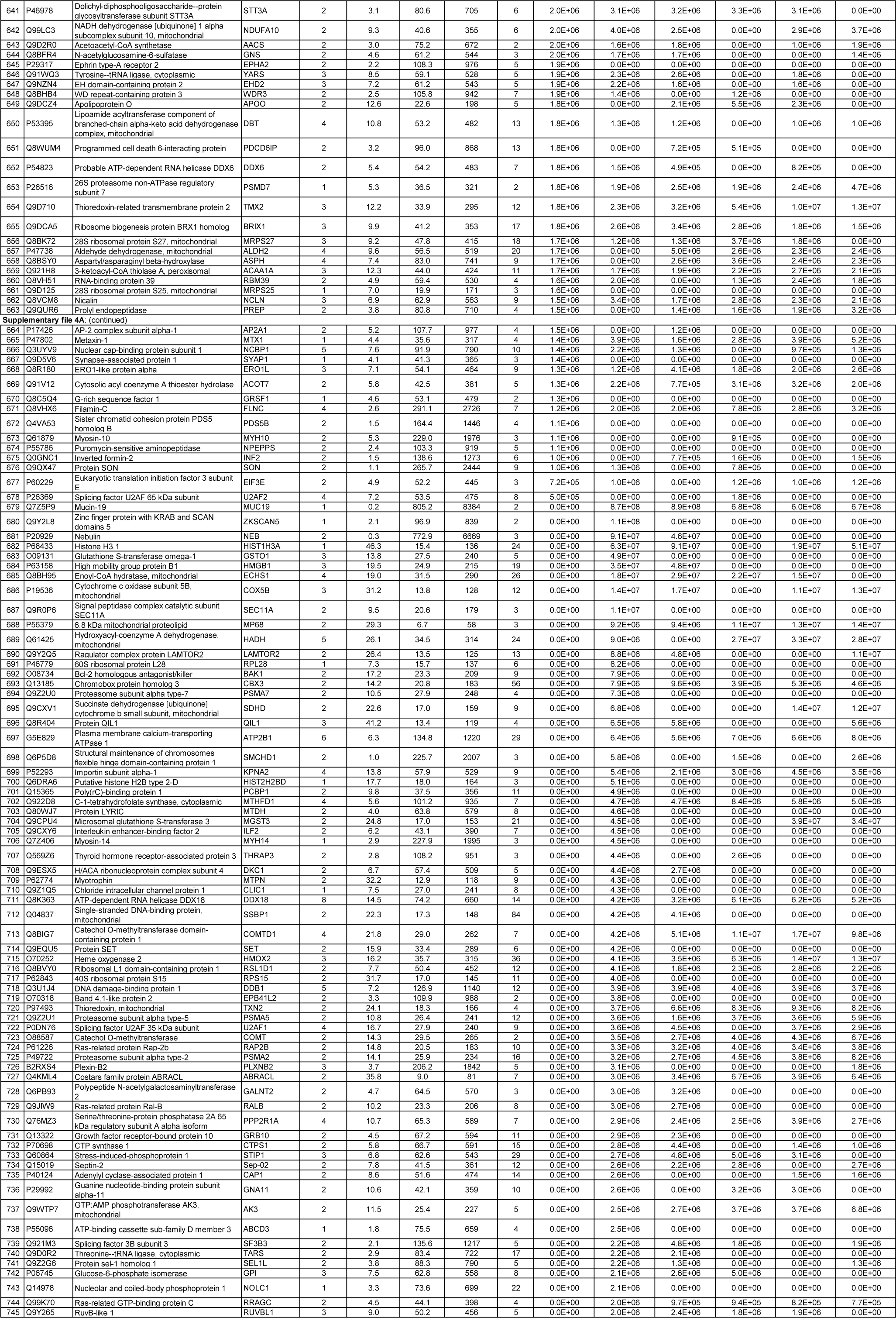
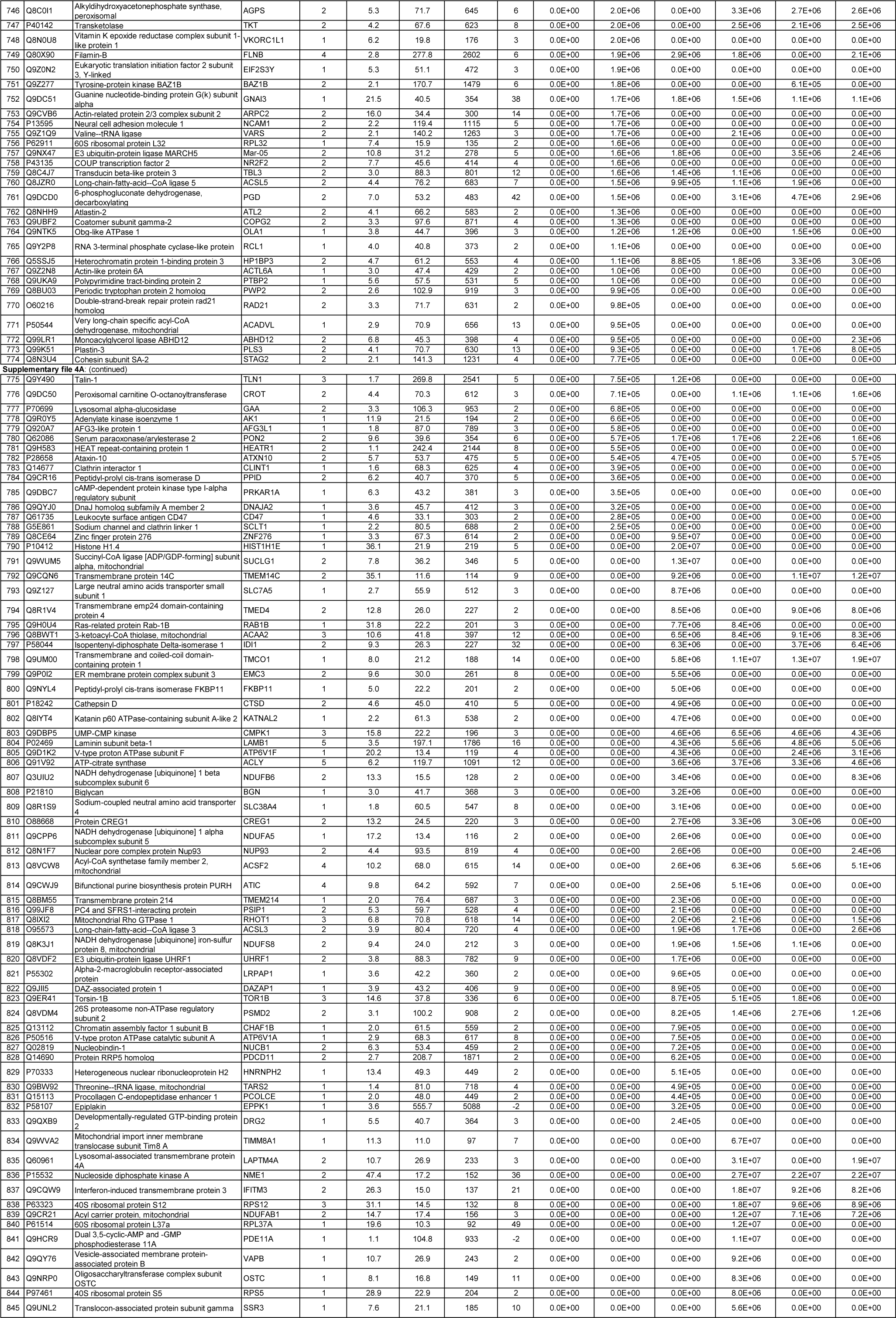
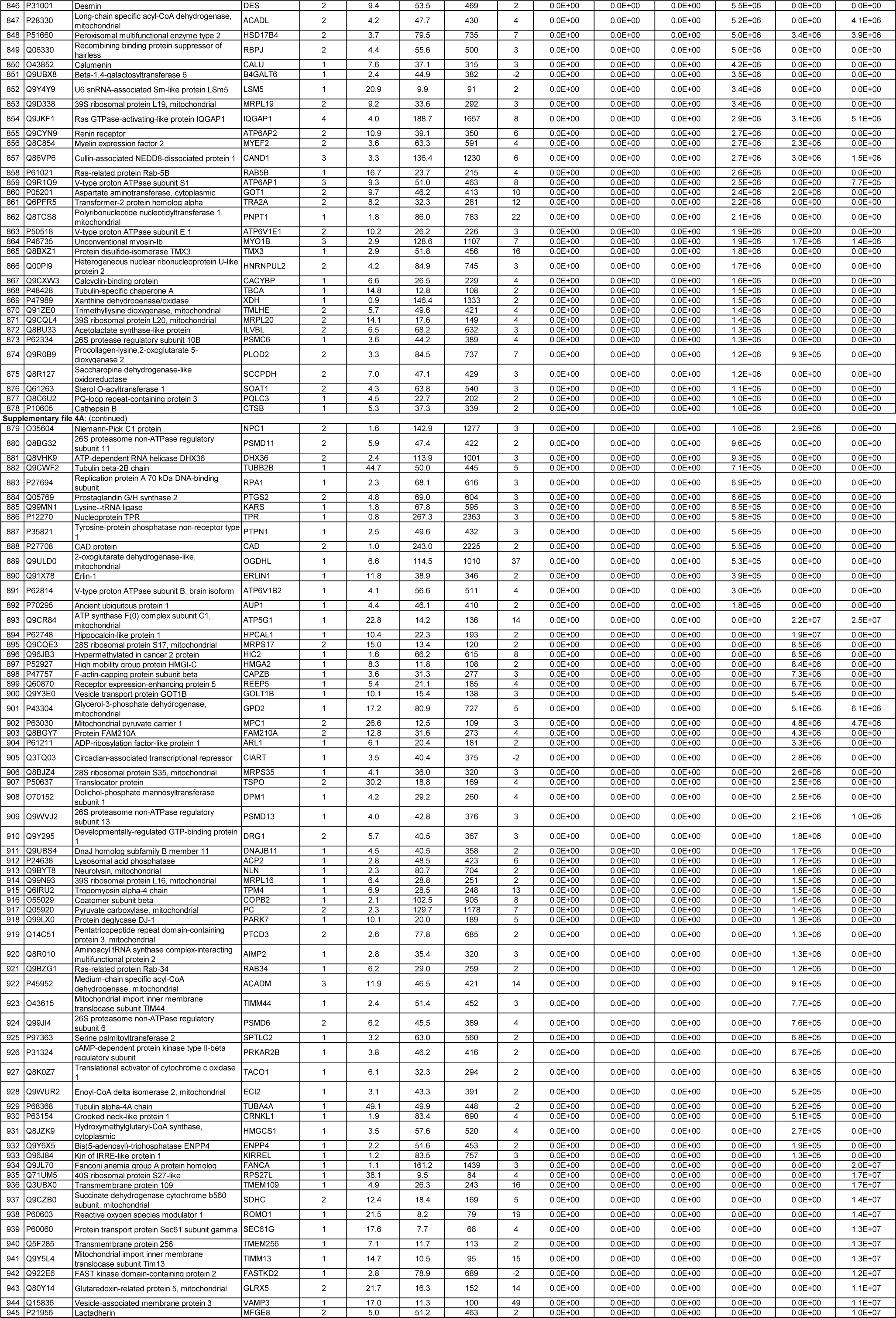
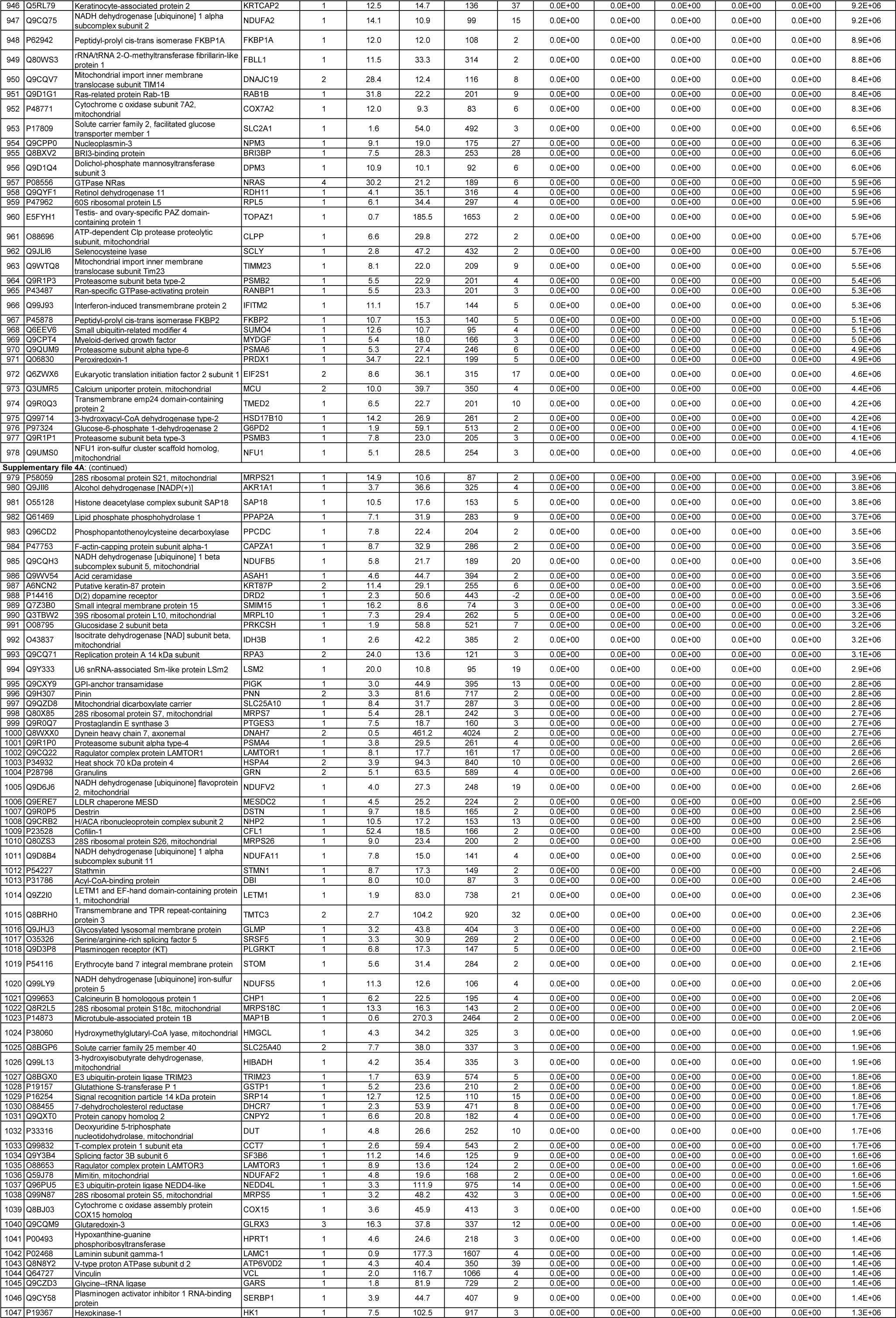
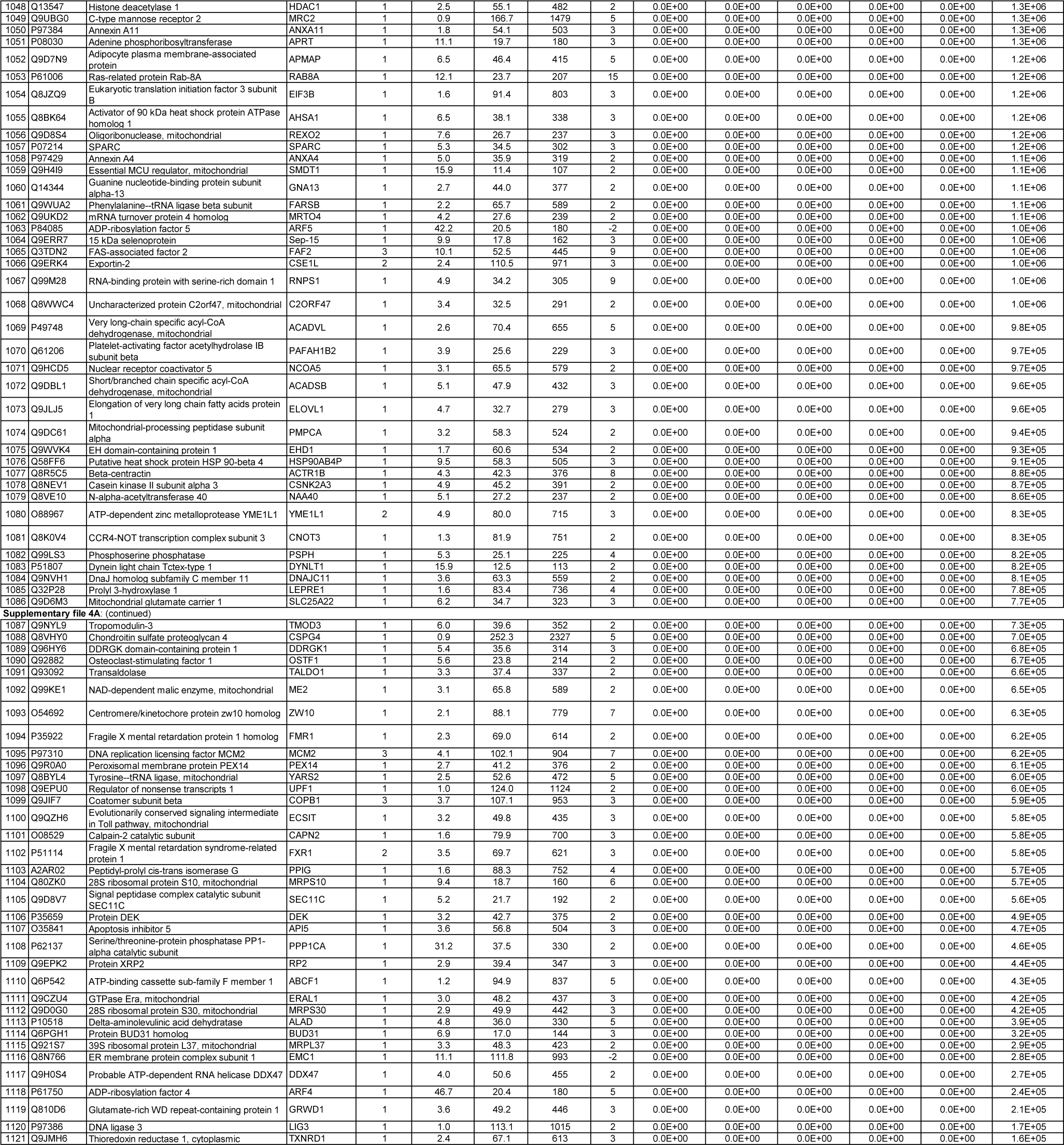
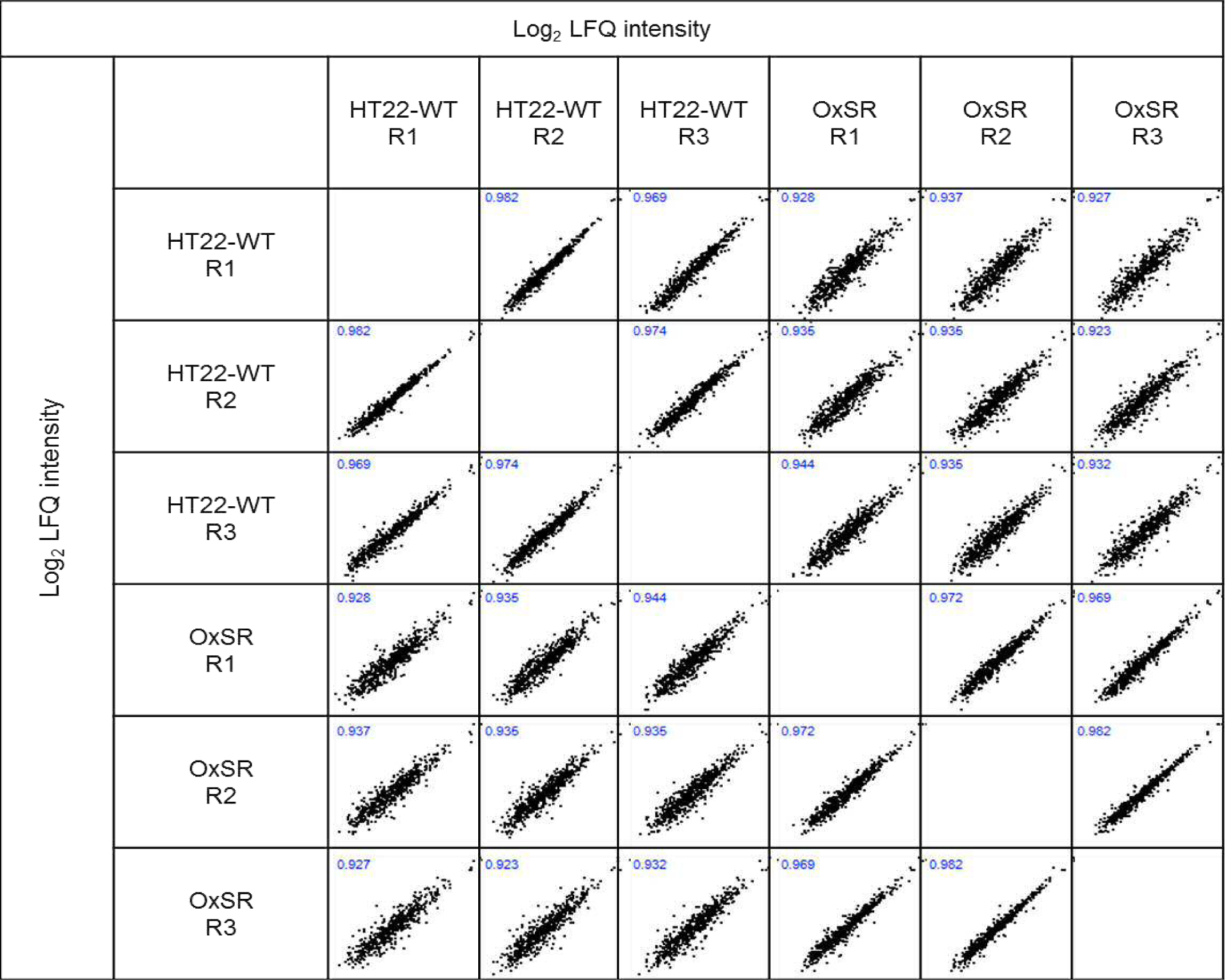
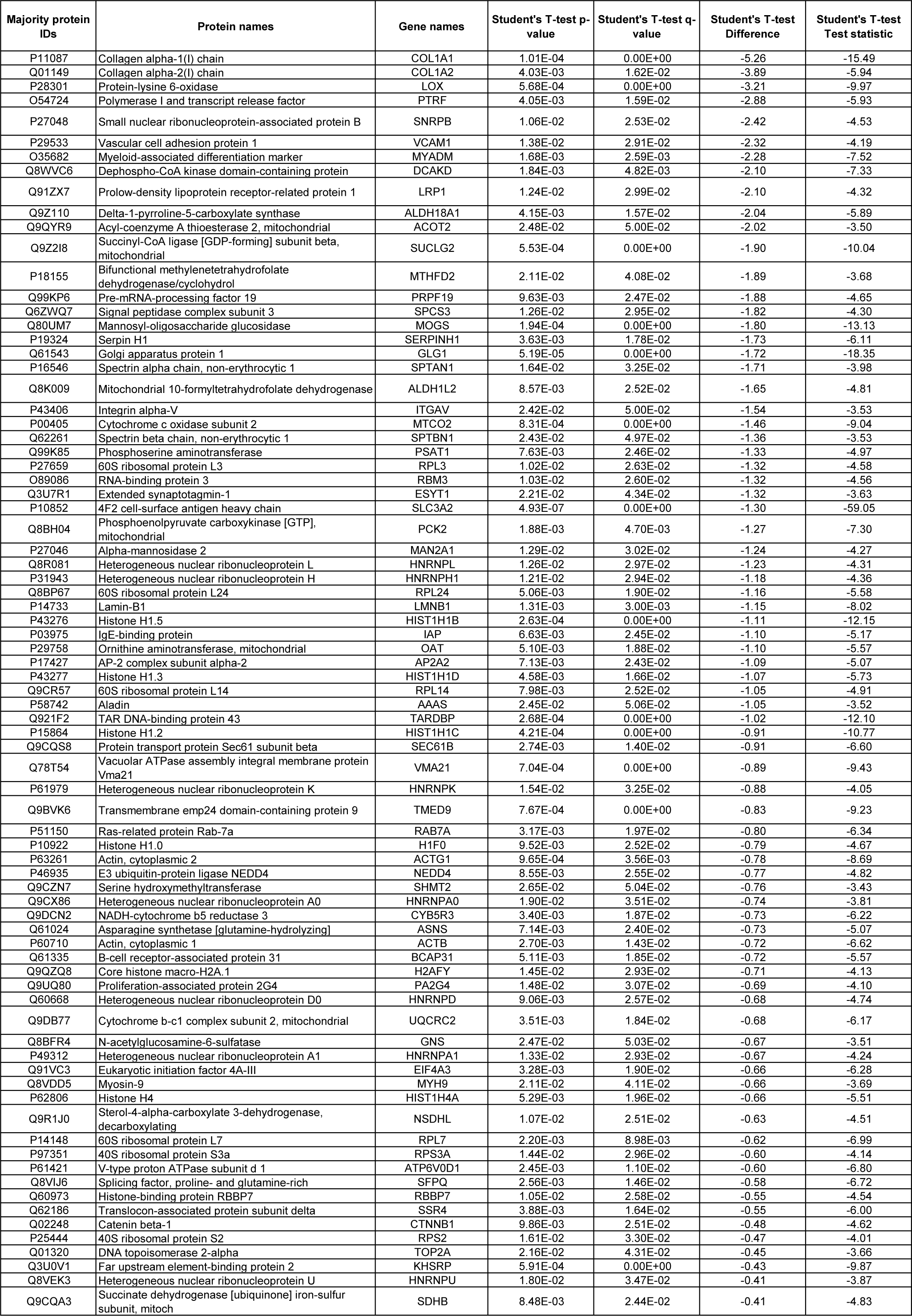
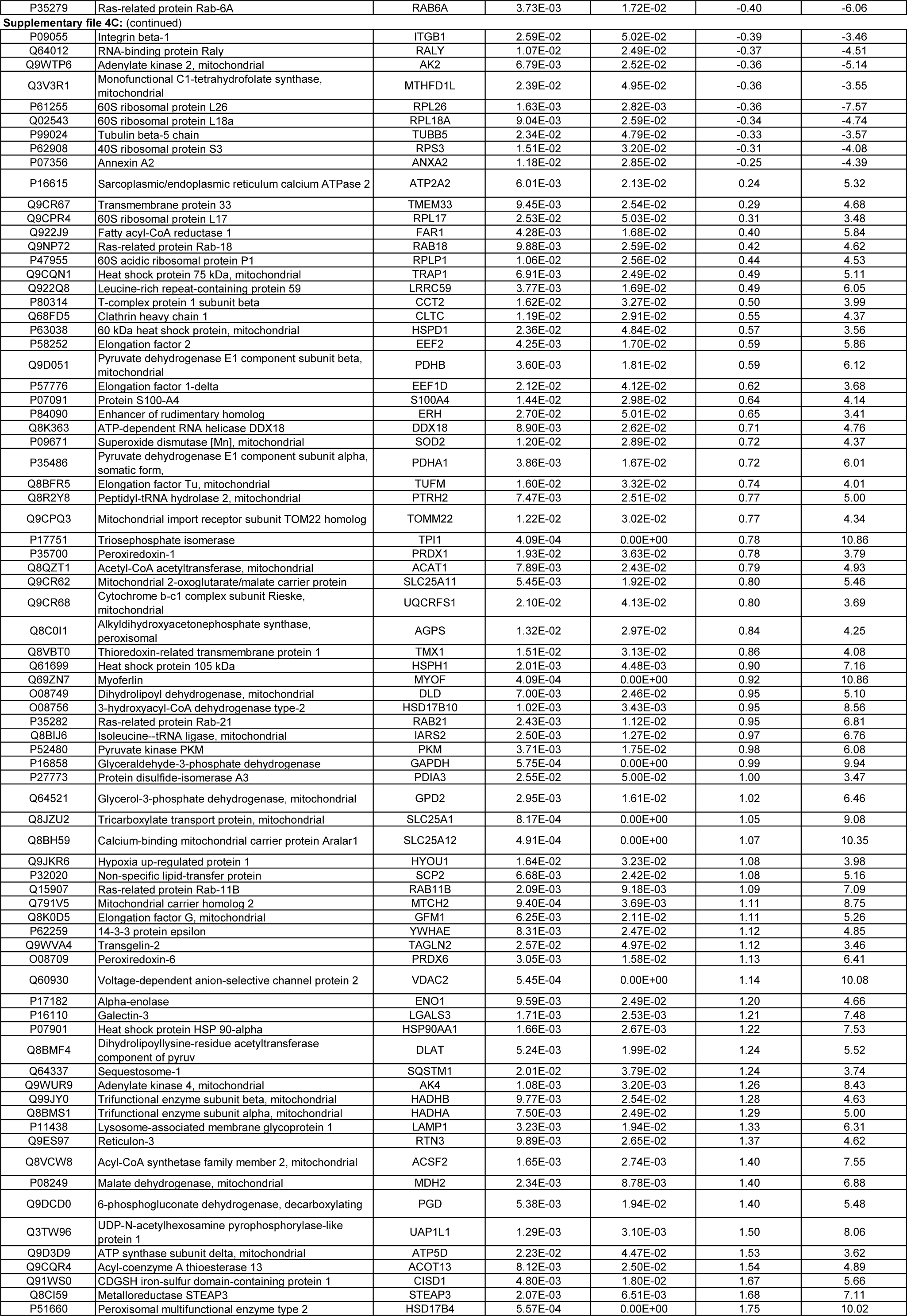
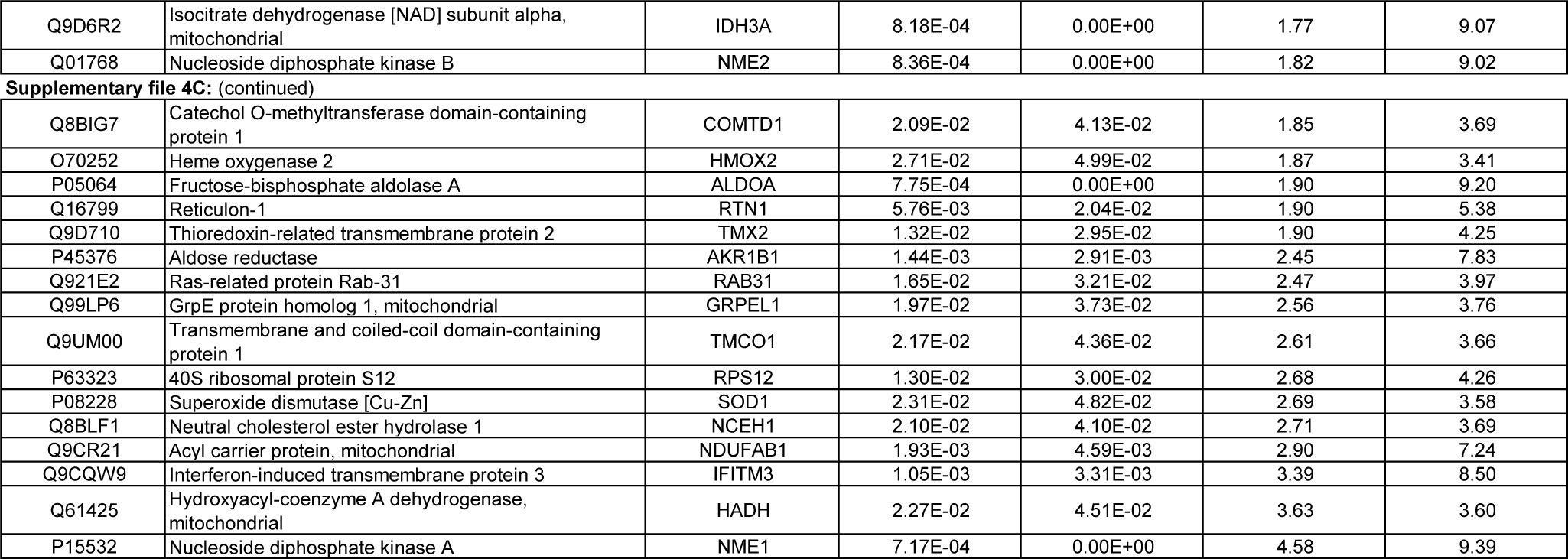
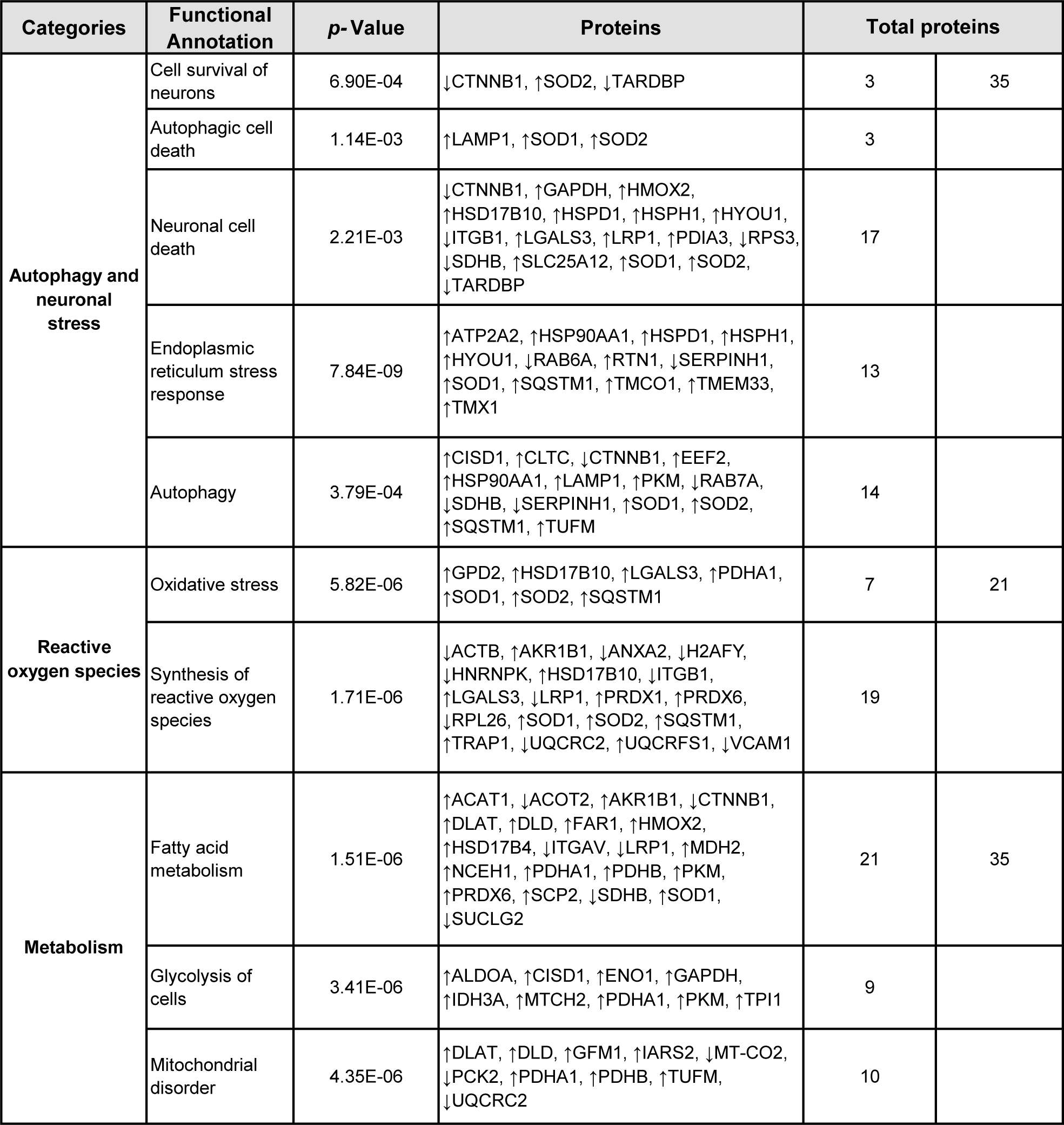
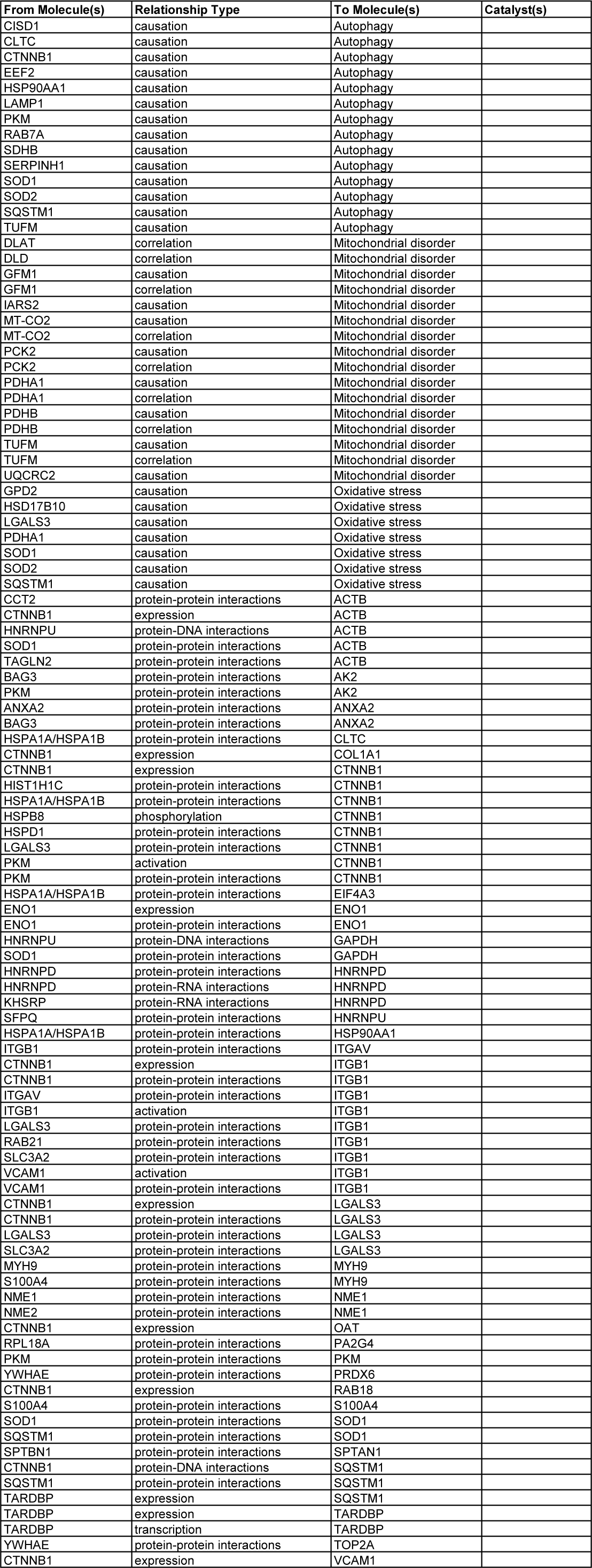
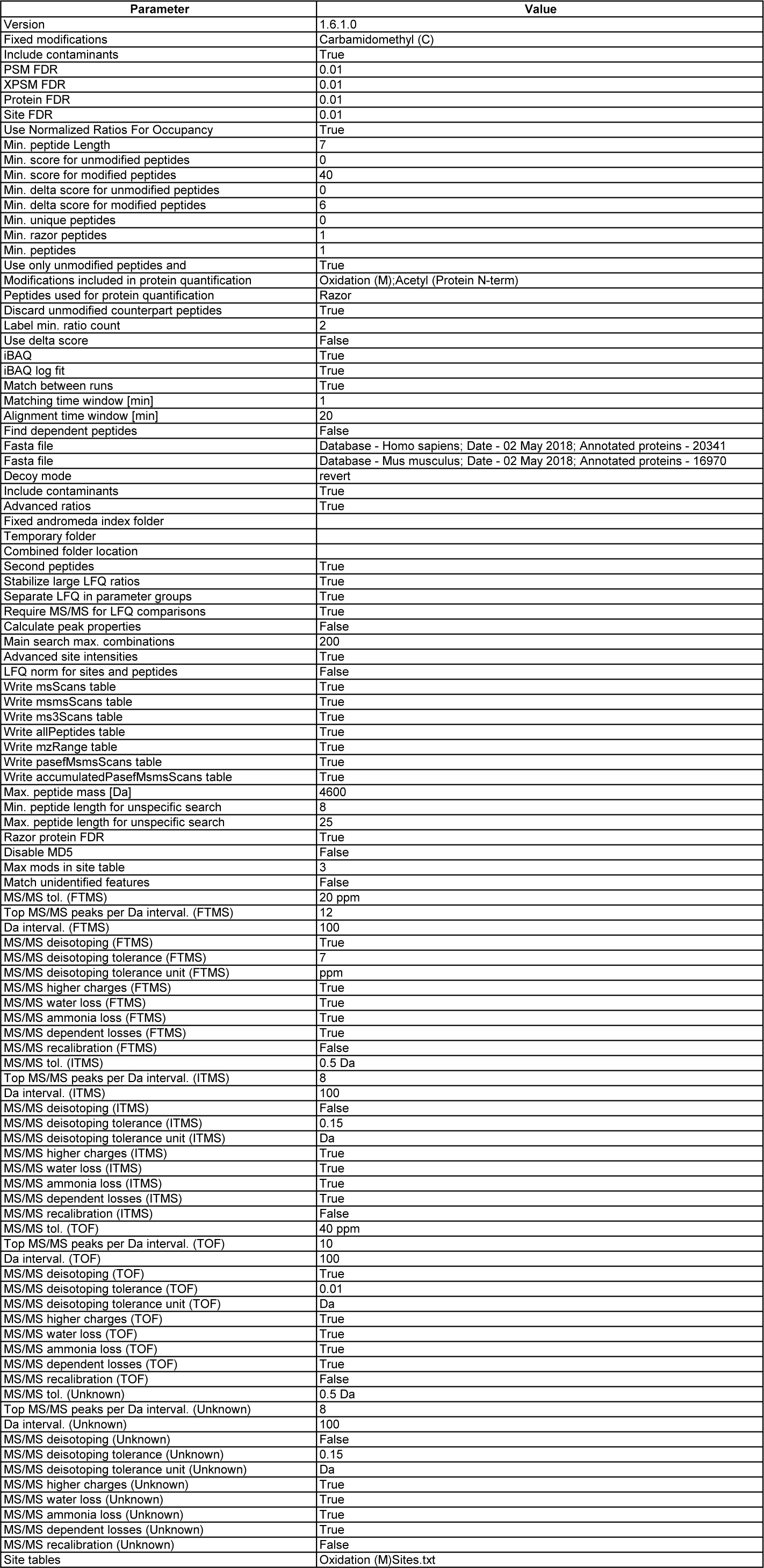
Proteomics data. (A) Complete list of all the proteins identified in proteome analysis of HT22-WT and OxSR cells employing MaxQuant. (B) The degree of variances in the proteome between the biological replicates and groups assessed by Pearson’s correlation analysis (C) List of the differentially expressed proteins in OxSR cells compared to HT22-WT cell line. (D) The detailed list of differentially expressed proteins in OxSR cells associated with each group of different cellular functions as indicated in Figure 5B. Upregulation and downregulation of expression in indicated by up (↑) and down (↓) arrows respectively. (E) Complete Protein-protein interactions network of the differentially expressed proteins in OxSR cells compared to HT22-WT cells as shown in Figure S2. (F) MaxQuant Parameters.

## Supplementary File S3 proteomics method

### Protein extraction

Protein extraction from the cell culture samples was carried out employing T-PER Tissue Protein Extraction Reagent (Thermo Scientific Inc., Waltham, MA, USA) according to an in-house established method catered especially for small amounts of samples, as described elsewhere [1]. Three biological replicates were utilized for each cell culture sample for this proteomics analysis. Briefly, 400 µl T-PER reagent was added to 70 µl samples from each replicate. Next, the samples were subjected to homogenization employing zirconium oxide beads in a bullet blender homogenizer (Bullet Blender Storm BBY24M, Next Advance Inc., Averill Park, NY, USA). Homogenates were centrifuged at 10,000 × g for 5 minutes to pellet tissue debris and, the supernatant was subjected to sample cleaning and buffer exchange employing the Amicon Ultra-0.5 centrifugal filter devices with 3K cutoff (Merck Millipore, Carrigtwohill, Ireland). Sample protein concentration of the obtained concentrate was determined using the standard bicinchoninic acid (BCA) Protein Assay Kit (Pierce, Rockford, IL) according to the manufacturer’s instruction.

### Sample preparation and 1-dimensional gel electrophoresis (1DE)

The samples were then subjected to 1DE (50 μg per well) employing precast NuPAGE 4-12% Bis-Tris 10-well mini protein gels (Invitrogen, Karlsruhe, Germany) with 2-[N-morpholino]ethanesulfonic acid (MES) running buffer under reducing conditions at a constant voltage of 150V in 4 °C. Pre-stained protein standard, SeeBlue Plus 2 (Invitrogen, Karlsruhe, Germany) was used as a molecular mass marker and the gels were stained with Colloidal Blue Staining Kit (Invitrogen, Karlsruhe, Germany), as per manufacturer’s instructions. Protein bands were excised (20 bands per replicate), reduced and alkylated prior to in-gel trypsin digestion employing sequence grade-modified trypsin (Promega, Madison, USA), as described in detail elsewhere [1]. Peptides extracted from trypsin digestion were purified using SOLAµ SPE HRP plates (Thermo Fisher Scientific, Rockford, USA) according to the manufacturer’s instructions. The resulting combined peptide eluate was concentrated to dryness in a centrifugal vacuum evaporator and dissolved in 10 μl of 0.1% trifluoroacetic acid (TFA) for LC-MS/MS analysis.

### Discovery proteomics strategy

Peptide fractionation was conducted in the liquid chromatography (LC) system, which consisted of a Rheos Allegro pump (Thermo Scientific, Rockford, USA) and an HTS PAL autosampler (CTC Analytics AG, Zwingen, Switzerland) equipped with a BioBasic C18, 30 x 0.5 mm precolumn (Thermo Scientific, Rockford, USA) connected to a BioBasic C18, 150 x 0.5 mm analytical column (Thermo Scientific, Rockford, USA). Solvent A consisted of LC-MS grade water with 0.1 % (v/v) formic acid and solvent B was LC-MS grade acetonitrile with 0.1 % (v/v) formic acid. The gradient was run for 60 min per sample as follows: 0-35 min: 15-40 % B, 35-40 min: 40-60 % B, 40-45 min: 60-90 % B, 45-50 min: 90 % B, 50-53 min: 90-10 % B: 53-60 min: 10 % B. The continuum MS data were obtained on an ESI-LTQ Orbitrap XL-MS system (Thermo Scientific, Bremen, Germany). The general parameters of the instrument were set as follows: positive ion electrospray ionization mode, a spray voltage of 2.15 KV and a heated capillary temperature of 220 °C. Data was acquired in an automatic dependent mode whereby, there was automatic acquisition switching between Orbitrap-MS and LTQ MS/MS. The Orbitrap resolution was 30000 at m/z 400 with survey full scan MS spectra ranging from an m/z of 300 to 1600. Target automatic gain control was set at 1.0 x 10^6^ ion. Internal recalibration employed polydimethlycyclosiloxane (PCM) at m/z 445.120025 ions in real time and the lock mass option was enabled in MS mode [2]. Tandem data was obtained by selecting top five most intense precursor ions and subjected them for further fragmentation by collision-induced dissociation. The normalized collision energy was set to 35 % with activation time of 30 ms with repeat count of 3 and dynamic exclusion duration of 600 s. The resulting fragmented ions were recorded in the LTQ. The acquired continuum MS spectra were analyzed by MaxQuant computational proteomics platform version 1.6.1.0 and its built-in Andromeda search engine for peptide and protein identification [3], [4], [5], [6], [7]. A target-decoy-based false discovery rate (FDR) for peptide and protein identification was set to 0.01. The summary of MaxQuant parameters employed in the current analyses is tabulated in Supplementary file S4F “Proteomics data”.

### Bioinformatics and functional annotation and pathways analyses

The output of the generated data from the MaxQuant analysis was utilized for subsequent statistical analysis with Perseus software (version 1.6.1.0). First, a log2 transformation of all LFQ intensities was done and the degree of variances in the proteome between the biological replicates and groups were assessed by Pearson’s correlation analysis [7]. Next, the data were filtered with minimum of three valid values in at least one group and the missing values were replaced from normal distribution (width, 0.3; down shift, 1.8) using the standard settings in Perseus. For statistical evaluation a Student’s two-sided t-test was used for the groups’ comparison with Permutation-based FDR ˂ 0.05 to identify the significantly differentially abundant proteins. Unsupervised hierarchical clustering analysis of the differentially expressed proteins was conducted with the z-scores of the LFQ intensities according to Euclidean distance (linkage = average; preprocess with k-means) and elucidated in a heat map. The list of the identified proteins was tabulated in Excel and their gene names were used for subsequent functional annotation and pathways analyses employing Ingenuity Pathway Analysis (v01-04, IPA; Ingenuity QIAGEN Redwood City, CA) (https://www.qiagenbioinformatics.com/products/ingenuity-pathway-analysis) [8]. IPA analyses elucidated the Gene Ontology Cellular Component (GOCC) terms, molecular types, protein-protein interaction (PPI) networks, and top disease functions associated with the proteins identified to be differentially expressed. Top canonical pathways of the differentially expressed proteins were presented with p-value calculated using Benjamini-Hochberg (B-H) multiple testing correction (–log B-H > 1.3). In PPI networks, proteins molecules are represented by their corresponding gene names and, only PPIs that were experimentally observed and had direct interactions were used.

